# Insights from U.S. beekeeper triage surveys following unusually high honey bee colony losses 2024-2025

**DOI:** 10.1101/2025.08.06.668930

**Authors:** Anthony Nearman, Christopher L. Crawford, M. Marta Guarna, Priyadarshini Chakrabarti, Katie Lee, Steven Cook, Elizabeth Hill, Arathi Seshadri, Garett Slater, Zac Lamas, Yan Ping Chen, Danielle Downey, Jay D. Evans

## Abstract

In January of 2025, U.S. commercial beekeepers reported unusually high honey bee colony losses as they prepared colonies for almond pollination. Two industry groups launched nationwide surveys to document colony losses between June 2024 and March 2025 across all scales of beekeeping. This study analyzes these survey data to assess colony losses, estimate financial impacts, and identify correlations with beekeeper management practices and geographical locations. Unlike past surveys, commercial beekeepers experienced more severe losses than smaller-scale beekeepers during this period. Respondents, managing over half of U.S. colonies, most frequently cited Varroa mites as the cause for their losses. Varroa mites were followed by pesticides and pathogens in the case of commercial beekeepers and by queen failure and weather in the case of smaller-scale beekeepers. Although Varroa was the most frequently cited cause, losses did not significantly differ between users and non-users of amitraz, suggesting that rising amitraz resistance alone does not explain observed trends. Differences in protein and carbohydrate feeding frequencies also played a role in net losses. While colony loss rates and financial concern varied widely among respondents, commercial beekeepers understandably showed higher sensitivity to financial impacts, with concerns increasing linearly with loss severity. This study highlights the value of beekeeper surveys which, alongside direct analyses of bee samples and longitudinal studies, help identify effective management strategies and environmental risks. Such insights are crucial for addressing the leading causes of colony losses on a national scale, and ultimately aid in safeguarding honey bee health, pollination services, and agricultural production.

**Highlights:**

- Unprecedented honey bee colony losses
- Indications of disease stress
- High economic pain for commercial beekeepers and growers

## 1 Introduction

Honey bees are vital players in agriculture and are important community members as they are both abundant in numbers and generalist pollinators. Across the globe, honey bees play an essential economic role for a wide variety of agriculturalists, ranging from small subsistence farmers to massive commercial operations with tens of thousands of colonies that add hundreds of billions of U.S. dollars to the worldwide economy (Khalifa et al. 2021, Stahlmann-Brown et al. 2023). In the United States, honey bees pollinate over 100 fruits, nuts, and vegetables, along with over 40 seed crops, sustaining billion-dollar crops in almonds and other high-value crops (Jordan et al. 2021). The role of honey bees as crop pollinators is at risk due to multiple biotic and abiotic stressors (Traynor 2015, Steinhauer et al. 2021, French et al. 2024), While the net number of honey bee colonies in the U.S. has remained constant for decades, high colony loss rates lead to costly efforts to protect and replace colonies.

Community-based surveys provide insights into annual changes in colony loss rates and can indicate which management practices and environmental stresses are correlated with differing colony survival success. The U.S. Beekeeping Survey has provided insights into the beekeeping industry and colony loss rates since its first iteration following heavy losses in 2006-2007. Funded by the USDA Animal and Plant Health Inspection Service and run first by the Bee Informed Partnership, and more recently by Auburn University and the Apiary Inspectors of America (Bruckner et al. 2023), these surveys show that beekeepers’ self-reported annual colony loss rates averaged approximately 40% over the past decade (Aurell et al. 2024). While these loss rates reflect averages, it has been apparent since the start of this survey that beekeeping losses are highly variable location-to-location, year-to-year, and beekeeper-to-beekeeper. In January 2025, U.S. commercial beekeepers notified the USDA-ARS Bee Research Laboratory of unusually high honey bee colony losses as they were preparing colonies to be moved for almond pollination. To investigate the scope and breadth of these claims, two industry organizations (Project *Apis m*. and the American Beekeeping Federation) developed and deployed community surveys to document colony losses, determine financial impacts, and correlate losses with management practices or geographical locations. This study reflects an analysis of the results of these two colony loss surveys.

By targeting beekeepers at all scales, these surveys generated data for over half of the managed bee colonies in the U.S. (Table 1). They recorded slightly larger losses than prior years over a nine-month window, along with tremendous variance across beekeeping operations in colony losses. Unlike prior years, commercial beekeepers (i.e., with >500 hives) faced especially high colony losses between June of 2024 and February of 2025, compared to sideliners (i.e., beekeepers managing between 50 and 500 hives) and hobbyist beekeepers (i.e., with <50 hives). We mined these losses for potential causes, including mite parasitism, pathogens, nutritional stress and pesticide exposure. Since commercial beekeepers have begun to adopt indoor ‘shed’ storage options as a way of increasing overwintering survival (Hopkins et al. 2021, Degrandi-Hoffman et al. 2023, Hopkins et al. 2023), we also explored outcomes for beekeepers using this method as opposed to outdoor storage.

*Varroa* mites and their associated viruses are often implicated in colony loss events (Dainat et al. 2012, Steinhauer et al. 2018, Stahlmann-Brown et al. 2022, Lamas et al. 2025). To investigate the roles of various *Varroa* mitigation strategies in recent losses, we correlated beekeeper reports on mite treatments against colony loss rates observed by the beekeeper. We focused especially on the use of the miticide amitraz, since *Varroa* mites have recently acquired widespread resistance in the U.S. to this common treatment (Hernández-Rodríguez et al. 2022, Rinkevich et al. 2023).

Additionally, we examined whether the type or frequency of carbohydrate and protein supplementation was correlated with colony losses. Supplemental feeding is a critical part of beekeeping management practices; this is especially true for beekeepers managing colonies during times of forage scarcity, which can be especially pronounced in some regions of the U.S. (DeGrandi-Hoffman et al. 2016, Chakrabarti et al. 2020, Tsuruda et al. 2021, Bernklau and Arathi 2023, Chakrabarti and Sagili 2023).

Loss rates were highly variable across operations, a result echoed by beekeepers’ financial sentiments in the past year. This mimics the pattern of loss events in 2023, wherein some beekeepers lost up to $1 million U.S. dollars due to economic hardships such as decreased revenue from unfulfilled pollination contracts and the costs of rebuilding colony numbers (Lamas et al. 2024). Commercial beekeepers were far more sensitive to the financial peril of their losses, showing a linear increase in financial concern. Sideliner beekeepers showed heightened concern when their losses topped 40-60% while hobbyists expressed significantly higher concern only when losses exceeded 80%. This survey demonstrates geographically widespread high colony losses that were similar across a range of operation sizes and management routines. As in prior reports, losses appear to reflect a range of factors ranging from parasitic mites and other biotic factors to environmental stresses.

## 2 Methods

### 2.1 Survey design and deployment

Two surveys were developed in parallel in early February 2025, when it was apparent that several commercial beekeepers in the U.S. were suffering extreme colony losses. One survey was driven by the industry nonprofit group Project *Apis m*. (P*Am*; https://www.projectapism.org/) and one by the industry trade group the American Beekeeping Federation (ABF; https://abfnet.org/). The P*Am* survey included 16 questions (Supplemental file S1) covering beekeeper operation size, geographical location(s), types of supplemental feeding, estimated ranges of fall mite levels, methods used to treat *Varroa* mites, queen replacement rates, methods of winter storage, perceived causes of colony loss, financial concern, and realized colony losses from June 2024 to February 2025. This survey was widely advertised in industry publications, online beekeeping groups, social media, and listservs dedicated to pollination and honey bees, resulting in ‘snowball sampling’ whereby early participants also recruited others. The survey ran from February 1, 2025, to March 15, 2025, capturing 842 unique responses (Table 1). P*Am* also distributed a second, follow-up survey to respondents who agreed to be contacted with additional questions. The purpose of the follow-up survey was to gather information on individual operation sizes (i.e., number of hives), which were in turn used to estimate the total number of colonies lost. This follow-up survey captured 110 unique responses, enabling the estimation of mean operation sizes for sideliners (mean = 314 hives, N = 6) and commercial beekeepers (mean = 6,798 hives, N = 90). When individual operation sizes were unavailable, these mean operation sizes were combined with individually reported loss estimates from the primary survey to estimate the number of colonies lost (Table 1; a value of five hives was assigned to hobbyists based on expert judgement, to provide a conservative estimate). The ABF survey presented similar questions regarding operation size, location, miticide treatments, queen replacement rates, winter storage practices, and realized colony losses from June 2024 to early 2025. This survey was sent to members of the American Beekeeping Federation, an industry group of 1000 members, and captured 107 unique responses (Table 1).

### 2.2 Data compilation and cleaning

Comma-delimited files with survey results were collected at the end of each survey period, stripped of personally identifying information and curated by scientists specializing in the various query topics. In both surveys, declared summer locations of the colonies were binned at the state level and subsequently grouped into U.S. Climate Zones, as delineated by NOAA (Karl and Koss 1984). When respondents managed colonies across multiple states, responses were attributed to the first state mentioned. Since several of the queries were open-ended, results were cleaned and binned as described in the Supplemental Methods. Beekeepers were classified according to operation size (i.e., number of hives) as follows: commercial (> 500 hives), sideliners (between 50-500 hives), and hobbyists (<50 hives). Cleaned datasets are made available as Supplementary Material, along with metadata describing each data field and the methods used to derive structured fields from the original survey questions. Metadata and details describing the data and associated cleaning and processing steps can be found in the Supplementary Information and Tables S4-S6.

### 2.3 Statistical analysis

All analysis was carried out using R programming language v4.5.0 “How About a Twenty-Six” (Team 2025). For each survey question, responses were categorized as binary or binned into multivariate categories either by biological relevance or statistically relevant sample sizes. Each question was then weighed against a set of common climate covariates. To achieve this, climate data was collected from PRISM (Daly et al. 1997) that represented the mean daily precipitation, total precipitation, and mean temperature from June 2024 through September 2024, as well as the mean minimum temperature from November 2024 through January 2025, for each U.S. state in the lower 48 and merged with the larger data set. This set of covariates was used to weight observations separately for each class of beekeeper (Commercial, Sideliner, and Hobbyist) using *WeightIt* v1.4.0 (Greifer 2019), with either the “ebal” (Hainmueller 2012, Zhao and Percival 2017) or “energy” (Huling and Mak 2024) method where required to achieve a balance threshold below 0.05 for all covariates. Rarely, classes of beekeepers who could not be balanced within a survey question, typically due to low response rate, were dropped from further significance testing, as indicated in the figures. As raw colony count data was available for only a subset of respondents, the reported proportion of colonies lost was incorporated as the response variable, along with the calculated covariate-balanced weights, into each individually modeled independent variable. Models were applied to a beta regression using *betareg* v3.2-2 (Cribari-Neto and Zeileis 2010). For multivariate questions, multiple comparisons were performed with *emmeans* v1.11.1 (Lenth et al. 2018) and compact letter display was mapped with *multcomp* v1.4-28 (Hothorn et al. 2012). All plots were generated using *ggplot2* v3.5.2 (Kassambara et al.).

## 3 Results

The P*Am* survey included responses from 280 commercial beekeepers (> 500 hives) and a total of 842 responses. Participation was balanced between small-scale beekeepers (“hobbyists” with <50 hives, N = 393), “sideliners” (50-500 hives, N = 161), and larger commercial beekeepers (Table 1). The ABF survey tallied responses from 87 commercial beekeepers (> 500 hives) and 20 smaller-scale beekeepers (< 500 hives, i.e., hobbyists and sideliners).

**Table 1.** Mean net colony loss rates captured in the two industry beekeeper surveys, conducted by Project *Apis m*. (P*Am*) and the American Beekeeping Federation (ABF). Results are shown by beekeeper class (i.e., operation size, in terms of number of hives), and across all respondents (“*Total*”). Mean values were calculated per respondent and represent the net loss rate experienced by the mean beekeeper. *Estimated number of colonies lost was calculated using data from P*Am’s* follow-up survey (see Methods).

| **Survey** | **Beekeeper Class** | **Mean Net Loss Rate** | **Number of responses** | **Estimated Number of Colonies Lost\*** |
| --- | --- | --- | --- | --- |
| **PAm** | Hobbyist (1-49 hives) | 51.2% | 393 | 974 |
|  | Sideliner (50-500 hives) | 53.9% | 161 | 26,645 |
|  | Commercial (>500 hives) | 62.3% | 280 | 1,147,613 |
|  | NA | 40.0% | 8 | - |
|  | *Total* | *55.3%* | *842* | *1,175,232* |
| **ABF** | Hobbyists and Sideliners (<500 hives) | 32.7% | 20 |  |
|  | Commercial (>500 hives) | 41.9% | 87 |  |
|  | *Total* | *40.2%* | *107* |  |

To ensure the quality of our covariates, we modeled each as independent variables against either Net Loss, Summer Loss, or Winter Loss as dependent variables. The only exception being mean minimum winter temperature, which was only modeled against Winter Loss. Significant results are presented in Table S1 and indicate our selected weather covariates are significant for predictors for Hobbyist and Sideliner beekeepers but not Commercial beekeepers. The only exception to this being mean daily precipitation during the active season, which was not significant for all classes of beekeepers for any period of losses. Significant associations with covariates were consistent between Sideliners and Hobbyists. Increasing mean minimum winter temperature was associated with lower winter losses (Figure S1). Similarly, increasing total active season precipitation and increasing mean daily active season temperature were associated with decreased net loss (Figure S2-S3). We also investigated the relationship between summer and winter loss reported by beekeepers and found that summer loss is a significant predictor for winter loss among all classes of beekeepers (Figure S4).

The P*Am* survey showed widespread losses across the country (Figure 1), while the ABF survey showed similarly widespread losses but at generally lower loss rates.

**Figure 1.**
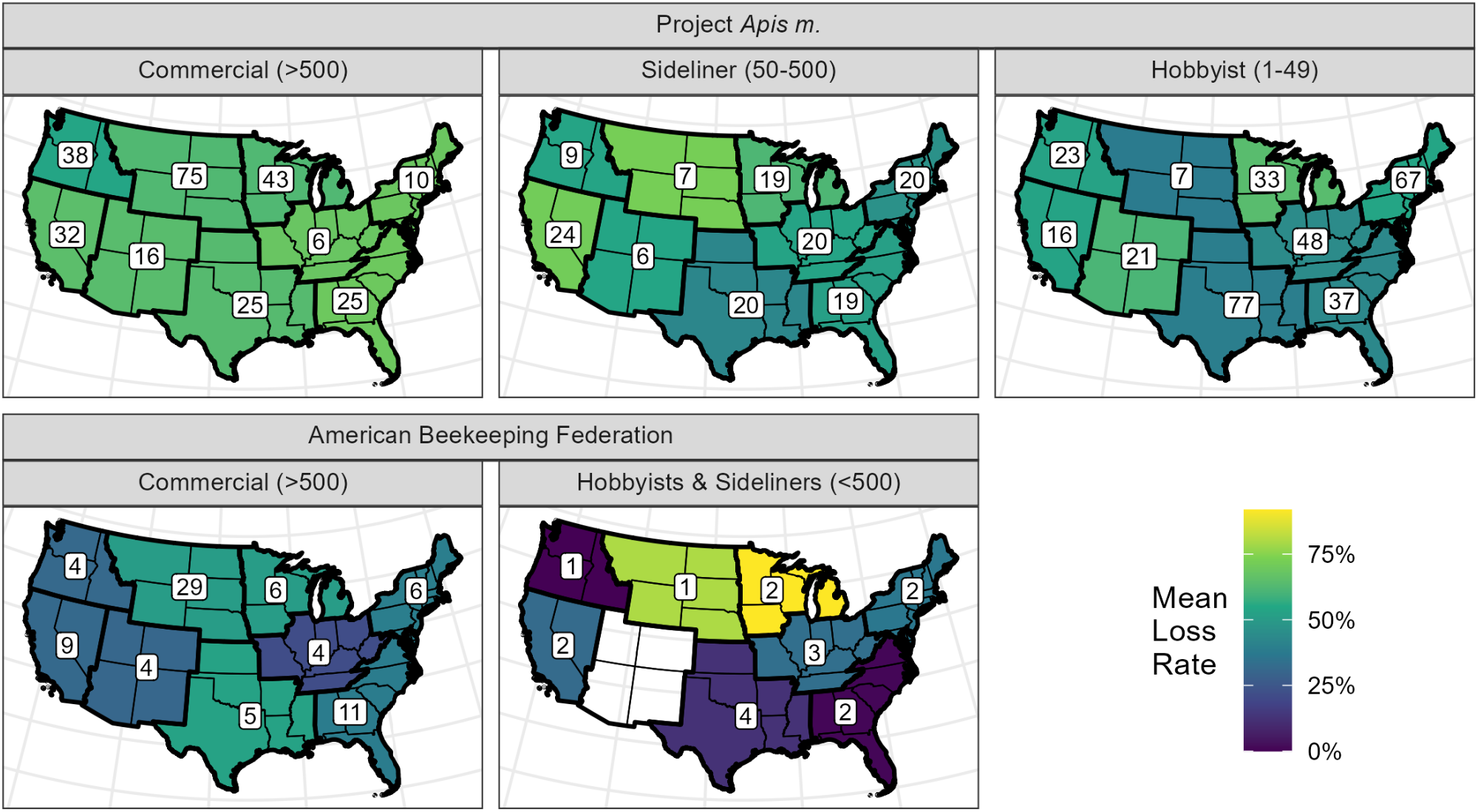
Mean net colony loss rates reported in two industry beekeeper surveys, shown by operation size (number of hives) and aggregated by region (U.S. Climate Zones, as delineated by NOAA; Karl and Koss 1984, [link](https://www.ncei.noaa.gov/access/monitoring/reference-maps/us-climate-regions)). Mean net loss rates are shown for beekeepers according to their first listed summer location and operation size according to the number of hives (noted in parentheses). Numbers of beekeepers in each group are shown in white boxes superimposed on each region.

**Figure 2.**
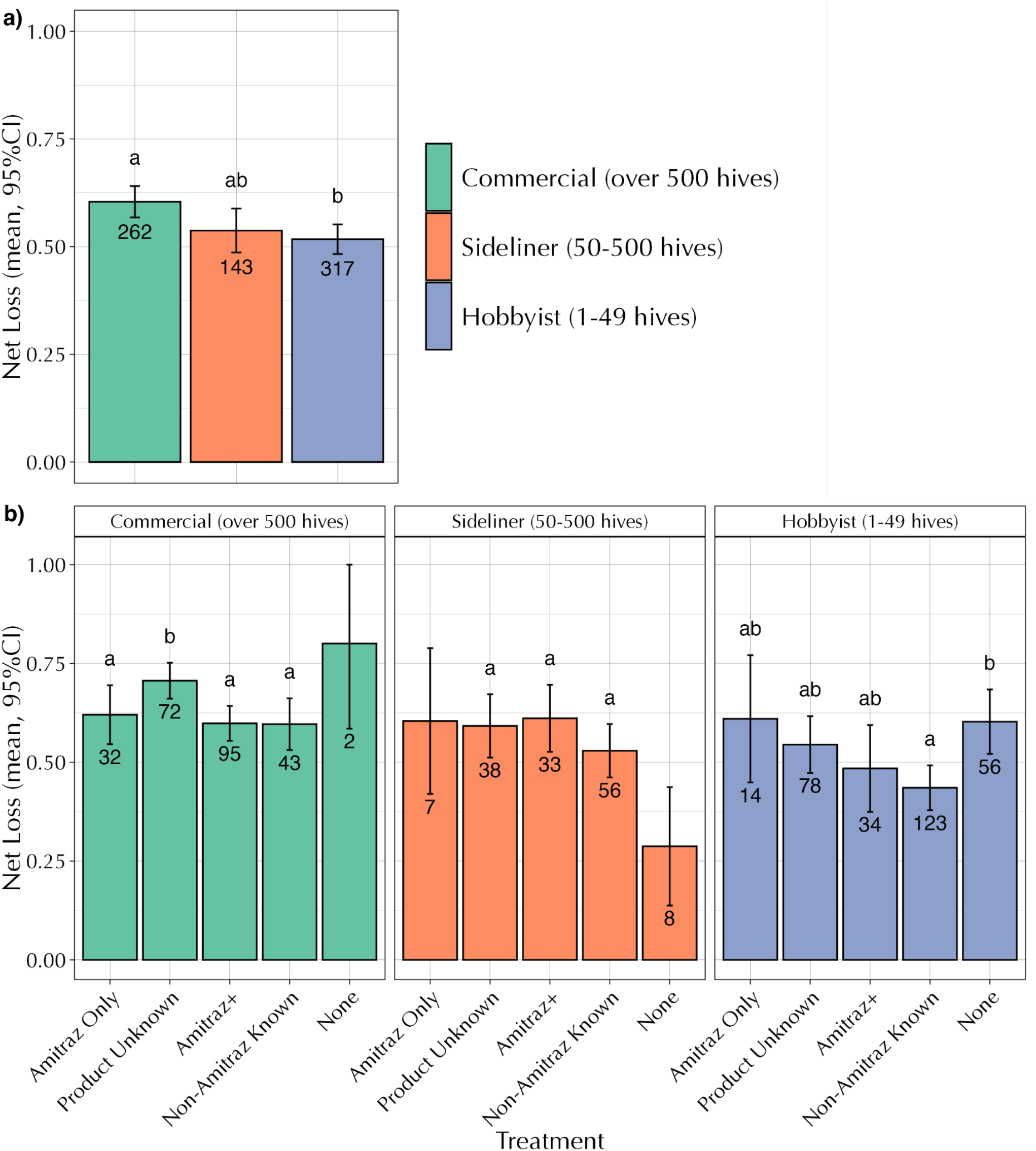
a) Colony loss rates by beekeeper class, P*Am* survey. b) Colony loss rates by miticide type, P*Am* survey.

Based on the P*Am* survey, commercial beekeepers suffered significantly higher losses than did hobbyist beekeepers (Figure 2a, *df*=2, χ^2^=11.925, Pr(>χ^2^)=0.003).

A total of 821 beekeepers provided information for the P*Am* question asking about product use and treatment frequency to control *Varroa* mites. Commercial beekeepers who did not identify the type of chemistry used as a miticide had significantly higher net colony losses than beekeepers who reported using only amitraz-based products, an amitraz-based product plus another chemistry, or used only non-amitraz based chemistries (Figure 2b, *df*=3, χ^2^=12.78, Pr(>χ^2^)=0.005). Beekeepers who reported using only non-amitraz products had similar losses to those who reported using amitraz (Figure 2b). Hobby beekeepers who did not use a chemical treatment had higher net colony losses compared to beekeepers who used non-amitraz based products (Figure 2b, *df*=4, χ^2^=13.336, Pr(>χ^2^)=0.010).

The most frequently used miticides among commercial beekeepers were amitraz and oxalic acid, reported by 133 and 115 of the 275 respondents to the P*Am* survey, respectively. For sideliners and hobbyists, oxalic acid was the most used treatment, reported by 85 of 160 sideliner respondents and 136 of 383 hobbyist respondents. The other commonly used miticides across all beekeeper groups were formic acid and thymol (Table S2). Beekeepers reported a treatment frequency range of 0 to 22 times in the June to December survey period. Commercial beekeepers reported treating their colonies an average of 4.7 times, compared to 3.5 times for sideliners and 2.1 times for hobbyists (Table S3). Of those who reported treating 20 or more times, one listed oxalic acid vapor, one rotated formic or thymol every 10 days, and the other three did not list the treatment used.

Similar results regarding amitraz use and colony losses were obtained from the ABF survey, even though the number of respondents was lower. Respondents to the ABF survey who used amitraz reported average losses comparable to those who did not use amitraz (Figure S5a). In addition, no differences in colony losses were observed between those who felt their mite treatments were effective and those who did not (Figure S5b). The most frequently used miticides were amitraz alone, and amitraz & oxalic acid (Figure S6), consistent with responses from the PAm survey. Beekeepers responding to the PAm survey were also asked to provide an estimated range of mite levels in their colonies. The most frequently reported range across all three beekeeper classes was 2-5 mites per 100 bees, although some did not test for Varroa levels. Analysis of these data revealed no clear differences in colony losses between beekeepers who provided different responses. One notable exception was among sideline beekeepers: those who reported not testing for Varroa experienced higher colony losses than those who monitored mite levels (Figure S7).

Supplemental feeding practices varied across regions, seasons, and beekeeper operation types (commercial, sideliner or hobbyist) as reported in the P*Am* beekeeper survey (Figure S8). For supplemental protein feeding, 231 commercial beekeepers responded “Yes” while 46 commercial beekeepers responded “No” to the question. For sideliner beekeepers, 95 responded “Yes” and 46 responded “No” to supplemental protein feeding. In the hobbyist beekeeper group, 140 respondents fed supplemental proteins to their colonies while 170 did not. There was no significant difference in net colony loss reported for commercial (Figure 3a; *df*=1, χ^2^=3.536, Pr(>χ^2^)=0.060), sideliner (Figure 3a; *df*=1, χ^2^=1.992, Pr(>χ^2^)=0.158) and hobbyist (Figure 3a; *df*=1, χ^2^=0.949, Pr(>χ^2^)=0.330) beekeeper groups with and without supplemental protein feeding. The commercial (Figure 3b; *df*=3, χ^2^=0.823, Pr(>χ^2^)=0.844) and hobbyist (Figure 3b; *df*=3, X2=4.462, Pr(> χ^2^)=0.216) beekeeper operation types did not significantly differ in their net colony losses when they fed their colonies protein supplements at different frequencies. However, the sideliner beekeepers reported a significantly higher net colony loss when they fed their colonies protein supplements only once (Figure 3b; *df*=3, χ^2^=11.960, Pr(> χ^2^)=0.008). Among beekeeper operations, 51 commercial beekeepers, 24 sideliner beekeepers and 47 hobbyist beekeepers fed their colonies proteins once during the year. Alternatively, 63 commercial beekeepers, 30 sideliner beekeepers and 33 hobbyist beekeepers fed their colonies proteins twice during the year, whereas 70 commercial beekeepers, 25 sideliner beekeepers and 34 hobbyist beekeepers supplemented their colonies with proteins more than three times during the year.

For the supplemental carbohydrate feeding survey question, 255 commercial beekeepers responded “Yes” while 6 commercial beekeepers responded “No” to providing their colonies with supplemental carbohydrates. For sideliner beekeepers, 127 responded “Yes” and 16 responded “No” to supplemental carbohydrate feeding. In the hobbyist beekeeper group, 258 respondents fed supplemental carbohydrates to their colonies while 55 did not. There was no significant difference in net colony loss reported for the sideliner (Figure 3c; *df*=1, χ^2^=0.258, Pr(>χ^2^)=0.611) and hobbyist (Figure 3c; *df*=1, χ^2^=0.001, Pr(>χ^2^)=0.982) beekeeper groups delineated by whether they did or did not feed sugar. As only six commercial beekeepers responded “no” to supplemental carbohydrate feeding, the weighted analysis was not balanced due to the low sample size (Figure 3c). Altogether, 122 commercial beekeepers, 81 sideliner beekeepers and 170 hobbyist beekeepers fed their colonies carbohydrates less than four times whereas 111 commercial beekeepers, 37 sideliner beekeepers and 67 hobbyist beekeepers supplemented their colonies with carbohydrates more than four times during this time period. Carbohydrate feeding frequency did not significantly affect loss rates for commercial (Figure 3d; *df*=1, χ^2^=1.045, Pr(>χ^2^)=0.307) or hobbyist (Figure 3d; *df*=1, χ^2^=0.697, Pr(>χ^2^)=0.404) operations. However, sideliner beekeepers experienced a significantly lower net colony loss when they fed their colonies carbohydrate supplements more than four times between June 2024 - March 2025 (Figure 3d; *df*=1, χ^2^=4.808, Pr(>χ^2^)=0.028).

**Figure 3.**
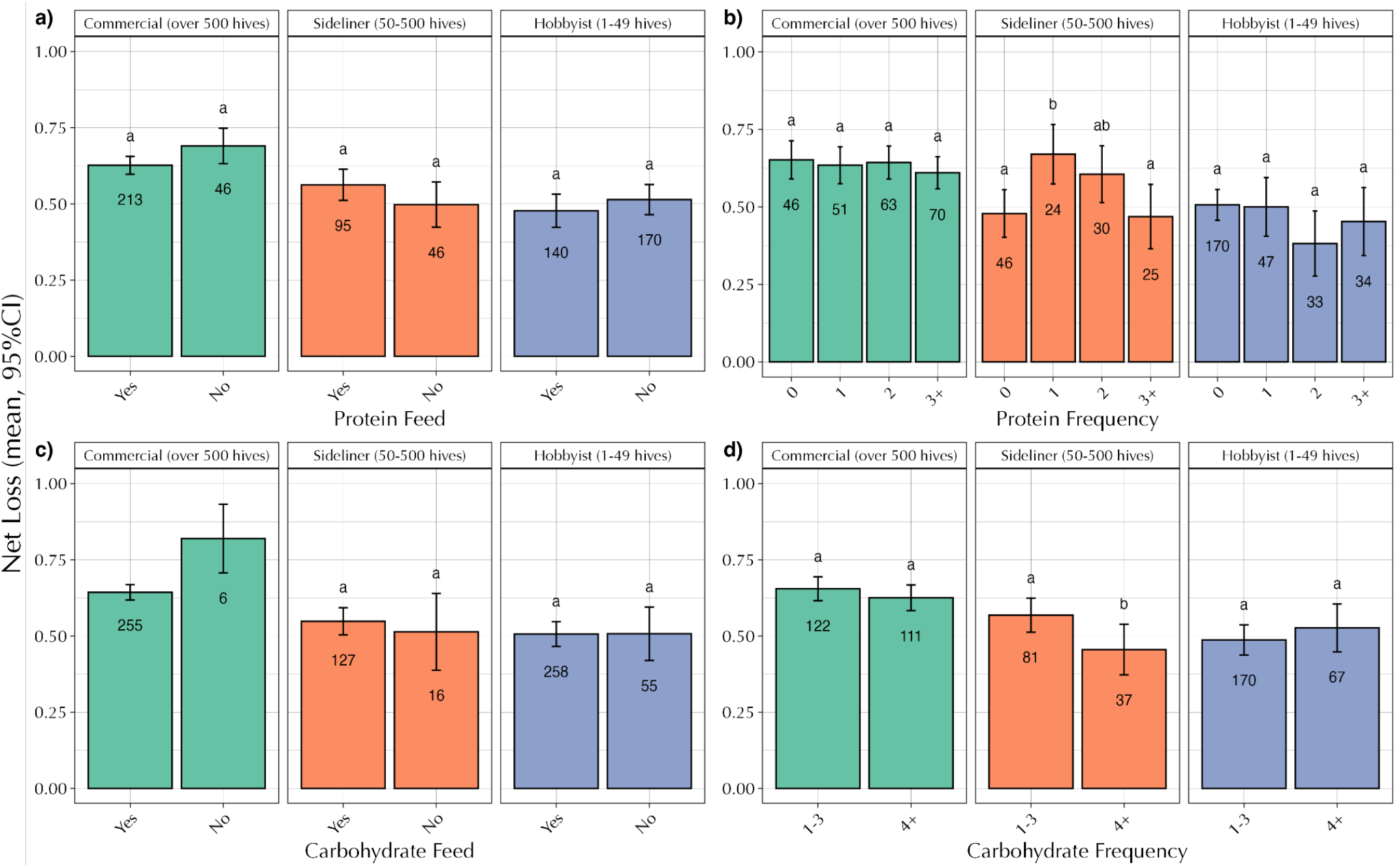
Weighted colony loss rates by supplemental carbohydrate or protein management as reported in the P*Am* beekeeper survey.

Other questions addressed management practices, including the number of queen replacements and whether colonies were overwintered indoors or outdoors. Analysis of these responses revealed that commercial operations reporting the highest percentages of queen replacement in the PAm survey also experienced higher colony losses (Figure S9). In contrast, similar losses were reported by beekeepers who used sheds for wintering their colonies and those who wintered outdoors. This pattern was observed among commercial beekeepers in the PAm survey, and among beekeepers with more than 50 colonies in the ABF survey (Figure S10).

**Figure 4.**
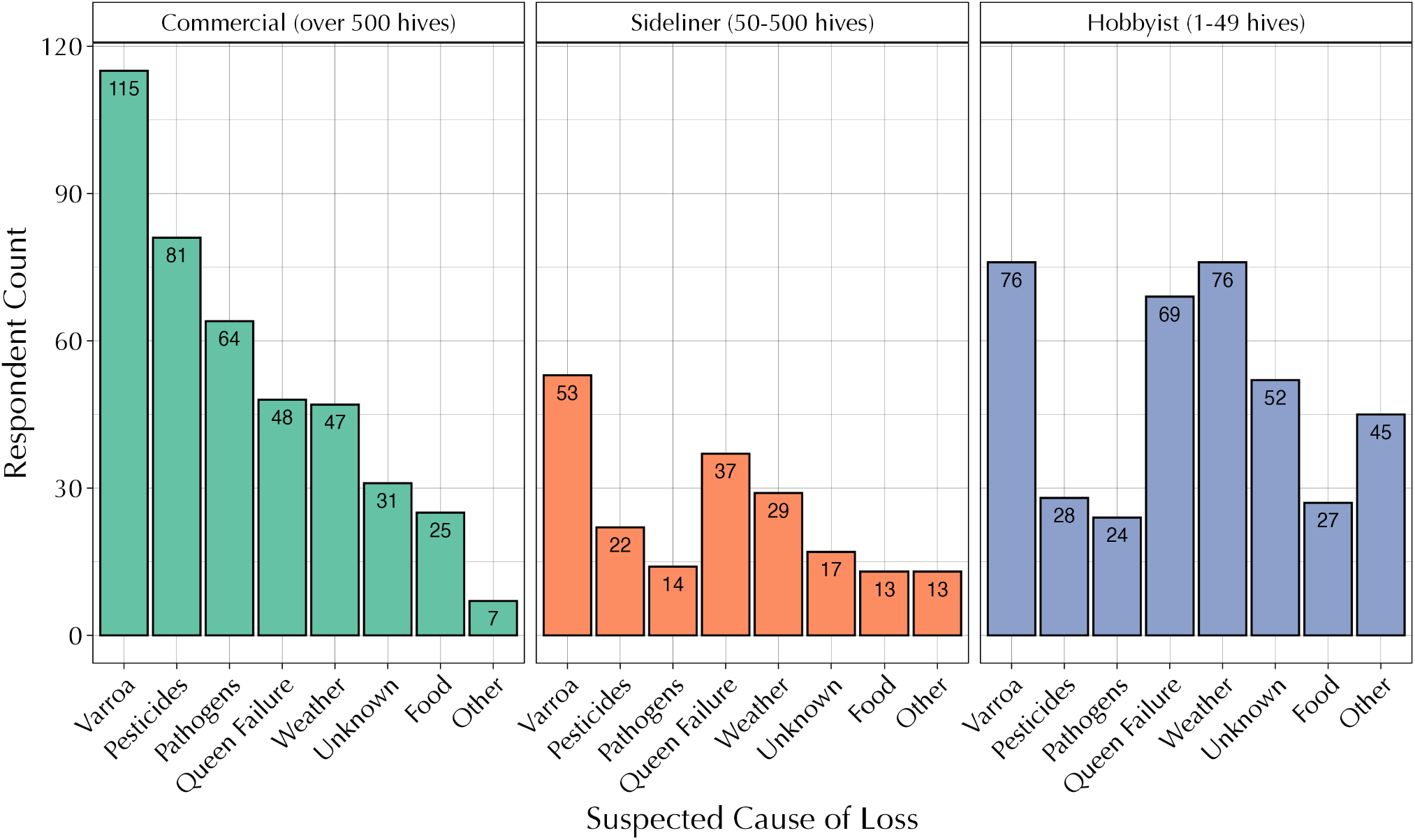
Suspected causes of losses reported by beekeepers in the P*Am* survey, shown by beekeeper class. Responding beekeepers could identify more than one possible cause of losses.

When prompted to report perceived causes of loss in the P*Am* survey, both commercial (n = 280) and sideliner beekeepers (n = 160) who chose a cause for their losses most frequently chose “*Varroa*” (Figure 4). For hobby beekeepers who chose a cause (n = 347), “*Varroa*” and “Weather” were chosen most frequently. Pesticides ranked second for commercial beekeepers, followed by pathogens (brood diseases and viruses that could be *Varroa* associated; of the 103 beekeepers who indicated pathogens, n = 27 indicated “viruses,” n = 78 indicated “disease,” and n = 2 chose both viruses and disease). Nine beekeepers wrote that losses were due to a hurricane. Beekeepers in the P*Am* survey could write in multiple stressors and frequently did. Notably, commercial beekeepers identified multiple perceived causes of colony losses more frequently than other beekeeper groups (18.9% compared to 12.5% for sideliners and 8.9% for hobby beekeepers).

**Figure 5.**
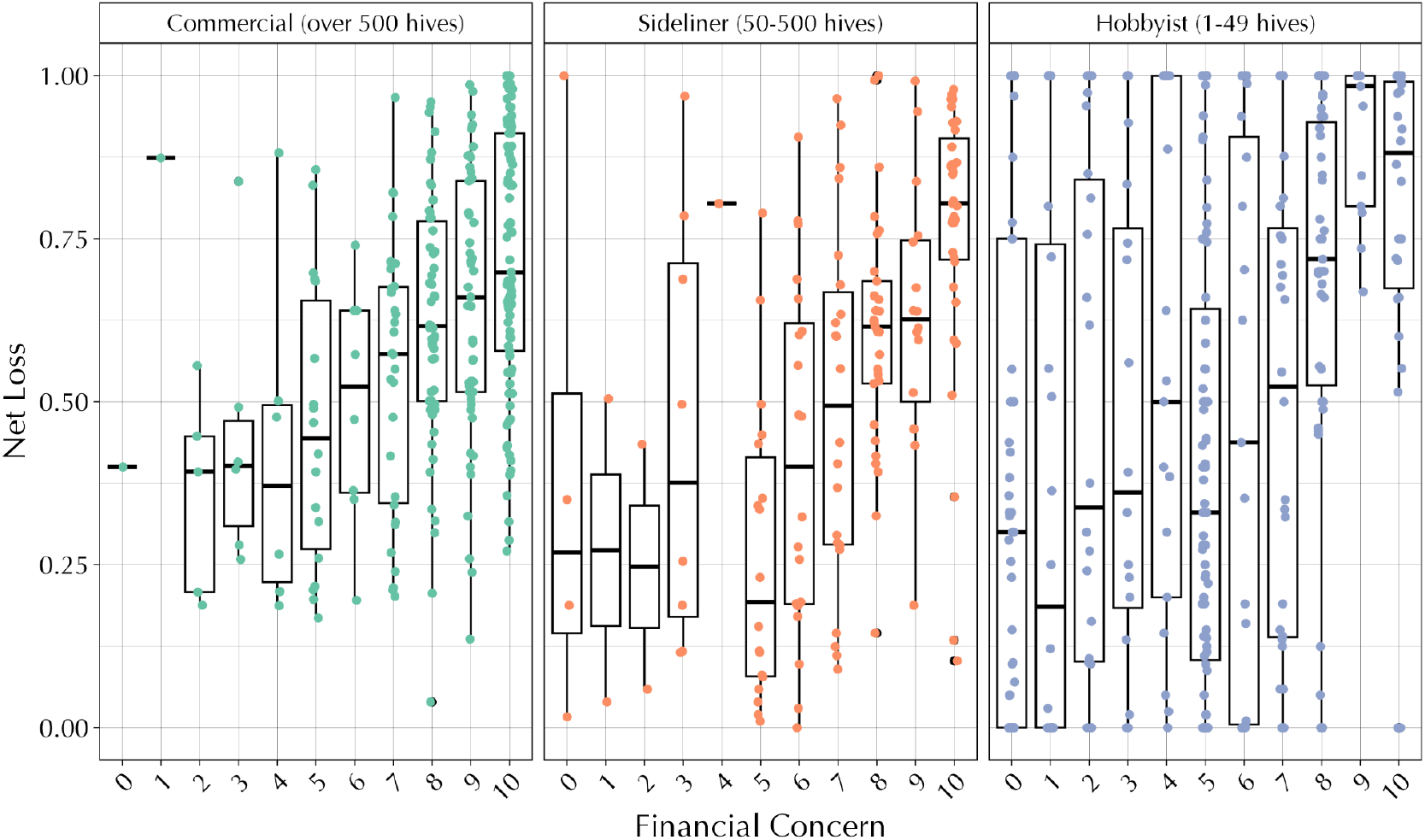
Financial concern of beekeepers compared to actualized losses, reported in the P*Am* survey.

Financial concern for commercial beekeepers trended highest across the three operation sizes, with concern levels rising in tandem with increasing colony losses (Figure 5). Thirty-two percent of the commercial beekeepers indicated that on a 10-point Likert scale, their financial concerns were a ten. One of the two commercial beekeepers who ranked financial concern as a 0, also put in “No, I quit” for the question about contacting them. A relationship between net loss and financial concern was also observed for hobbyists and sideliners, with sideliners tending to show increased concern as losses went above ca. 30% while hobbyists tended to express heightened concern only when losses exceeded 75%. The relationship between financial concern and loss was stronger for sideliners than for hobbyists. This makes sense, as sideliners rely on their colonies for income more than hobbyists do, though not as heavily as commercial beekeepers.

## 4 Discussion

Honey bees are critical to our agroecosystem but maintaining colonies year after year has become an increasing challenge. Beekeepers devote a large fraction of their budgets to the labor and material costs of rebuilding annual losses, reducing disease stress and providing supplemental feed. Despite these investments, beekeepers can lose entire operations suddenly, with devastating personal and industry-level economic impacts (Lamas et al. 2024). Traditional local knowledge has often been relied upon in a Multiple Evidence Based approach to inform decisions for management (Smith et al. 2017). Thus, beekeeper perceptions and their colony management history can often help in interpreting the reasons for colony decline and losses. Sudden colony loss events, as observed in 2024-2025, offer an opportunity to explore and strive to mitigate these potential causes. The two surveys described here, although not as complete as long-standing U.S. and worldwide colony health surveys (e.g., surveys conducted by the USDA National Agricultural Statistics Service), offer insights into the scope and key features of loss events. The U.S. bee losses of 2024-2025 are striking in that they were so widespread across the entire country (Figure 1). While losses were reported starting in the summer of 2024, many of the most severe losses occurred as bee colonies were pulled from winter storage in advance of profitable pollination events.

Even though there is evidence of miticide resistance to amitraz (Hernández-Rodríguez et al. 2022, Rinkevich et al. 2023), beekeepers who reported using other products in lieu of amitraz had similar losses to those beekeepers who used amitraz exclusively or in combination with another chemistry. In contrast, commercial beekeepers who identified a specific miticide product or products experienced significantly lower net colony losses than those who did not name the type of miticide used in their colonies. This suggests that colony losses may be influenced by the efficacy of the products used and/or *Varroa* mite resistance to amitraz. Although the initial analysis of reported fall 2024 mite levels didn’t reveal clear associations with colony losses, future investigations exploring associations between mite loads, mite mitigation strategies, and colony losses may be informative. Overall, these results highlight the complexities in interpreting the observed patterns and the need for field-vetted data.

Our choice to model weather characteristics as covariates was meant to provide higher resolution, in terms of local management practices and bee behaviors, compared to simple geographic location. For example, stationary apiaries in colder, northern climates have shorter active seasons and are exposed to fewer or different floral resources compared to their southern counterparts. Weather is also critical for honey bee success and survival (Calovi et al. 2021, Insolia et al. 2022, Overturf et al. 2022). By removing the effects of local climate, we were better positioned to estimate the effects of reported management decisions in the survey. In evaluating any direct associations between our weather covariates and reported loss, we found significant associations only among sideliners and hobbyists. This may be because these groups are variably stationary, or completely in the case of hobbyists, and more subject to local weather changes. Further, commercial operations may either be migratory or span multiple geographic locations, avoiding or diluting any weather-related losses.

Our work suggests climate variables are significant among presumably stationary and single-location sideliners and hobbyists. Recent work, however, indicates that regions commercial beekeepers rely upon for staving off the effects of dearth or climate may be experiencing extremes regarding drought, precipitation, or changes to the landscape tending towards being more crop-dominated (Morton et al. 2015, Ahlering et al. 2020, Hoell et al. 2021, Haigh et al. 2022, Heim Jr et al. 2023). While extreme drought and precipitation may have measurable effects on crop yields and pollinator friendly volatiles (Kuppler and Kotowska 2021) and products (Vaudo et al. 2024), the downstream effects on pollinator health appears to vary per location and pollinator, with some work indicating a range of tolerance, that may relate to changes in landscape (Descamps et al. 2021, Brunet et al. 2025). Our data suggest the locations and subsequent losses reported by commercial beekeepers are unrelated to climate variables, but should that change under future surveillance efforts, those data would be invaluable to identifying the locations and conditions that lead to increased losses for commercial beekeepers. As such, loss surveys should continue to monitor any relationships to climate-related variables and colony losses moving forward. Additional work may also be performed on the data published here to determine any location-specific effects among commercial beekeepers, though the data does not discern between migratory and multi-state beekeepers and their respective losses in each location.

We also found that reported summer losses predicted winter losses, an effect that was most prominent among commercial beekeepers (Figure S4). This appears to be driven by beekeepers who experience all around lower losses (<25%), rather than a one-to-one correlation. Still, the data indicate that there are beekeepers who experience variable rates of summer and winter loss, outside of only low or only high net losses. Additional analyses should investigate any relationships between management decisions and the variability of loss across all seasons.

Supplemental feeding is often an important part of beekeeping management and the beekeepers’ location, weather conditions, participating in pollination services and colony needs may drive the demands for supplemental protein and carbohydrate feeding throughout the year. Sideliner beekeepers reported a significantly higher net colony loss when they supplemented their colonies with proteins only once and carbohydrates less than four times annually, compared to other groups. This reiterates the need for further assessments of local forage availability, colony densities and weather conditions for colonies which may drive their dependence on supplemental feeding.

Beekeepers self-reporting causes associated with losses most frequently chose *Varroa*, consistent with past beekeeper loss surveys (Aurell et al. 2024). However, other contributors were also identified as being important, including pesticides, viruses, nutrition, or regional weather events. *Varroa* mite levels are immediately measurable, while contributions by other factors, like pesticides and nutrition, can be less apparent. In addition, colonies weakened by factors such as pesticides and poor nutrition may become susceptible to *Varroa* mite infestation, obscuring the primary cause.

While other surveys have measured high honey bee losses, including surveys conducted by the U.S. Department of Agriculture’s National Agricultural Statistics Service (NASS; https://www.nass.usda.gov/Surveys/Guide_to_NASS_Surveys/Bee_and_Honey/) and Auburn University, the described P*Am* and ABF surveys were critical in rapidly quantifying the scope of specific in-field issues experienced by beekeepers. Responses to the P*Am* survey included 280 commercial beekeepers, while the commercial responses to three recent years of the U.S. Beekeeping Survey averaged only 41 commercial beekeeper respondents (Bruckner et al. 2023). Self-reporting, optional surveys can be prone to response bias, and it is possible that the commercial beekeepers who responded to the P*Am* and ABF surveys were more likely to have experienced the higher loss rates. However, with over 50% of U.S. colonies represented by participants, higher than all prior surveys, the results are likely to strongly reflect the industry. These surveys demonstrate the financial strain faced by commercial beekeepers, as evidenced by 32% of all commercial beekeepers responding to the P*Am* survey ranking their financial concern at the highest level. These sociological indicators, coupled with in-field measurements of loss events (Lamas et al. 2024), demonstrate the precarious nature of bee pollination, a key agricultural service.

## Supporting information

ABF_Data

ABF_DataDescription

PAm_Data

PAm_DataDescription

ColonyLossSurvey_SupplementalMaterials

## 5.Acknowledgements

We are deeply grateful to the beekeepers who took the time to respond to these surveys. CLC was supported by an AAAS Science & Technology Policy Fellowship served at the USDA Office of the Chief Scientist.

## 6 Contributions

Conceptualization: Jay Evans, Danielle Downey

Data curation: Priyadarshini Chakrabarti, Steven Cook, Christopher L. Crawford, Katie Lee, Anthony Nearman

Formal analysis: Anthony Nearman, Christopher L. Crawford Funding acquisition: N/A

Investigation: Jay Evans, M. Marta Guarna, Danielle Downey

Methodology: Anthony Nearman, Christopher L. Crawford, Priyadarshini Chakrabarti, Katie Lee

Visualization: Christopher L. Crawford, Anthony Nearman

Writing – original draft: Jay Evans, Priyadarshini Chakrabarti, M. Marta Guarna, Katie Lee, Anthony Nearman

