## Supplementary material for "Insights from U.S. beekeeper triage surveys following unusually high honey bee colony losses 2024-2025": ColonyLossSurvey_SupplementalMaterials

### Supplemental Methods

#### Research Data:

The cleaned survey data collected separately by Project *Apis m.* (P*Am*) and the American Beekeeping Federation (ABF) are publicly archived and available via Zenodo (or other repository; formal reference forthcoming). Any data that could reveal the identities of individual respondents were removed. Data that contained personal identifiers were removed from columns including Percent_Almonds, Predicted_Cause, Pesticide_Details, Other_Pests, Supplemental_Feed_Sugar, Supplemental_Feed_Protein, Queen_Replacement, Percent_Indoor, Percent_Loss_Indoor_Outdoor, Loss_Summer_Fall_Clean, Loss_Summer_Fall_n, Loss_Winter_Clean, and Loss_Post_Emergence_Clean. A companion metadata file, listing the field (column) names and descriptions, including descriptions of how derived data fields were subsequently cleaned and categorized, is also archived together with the data. Additional data requests will be considered on a case-by-case basis.

#### Survey Data Processing:

The original P*Am* survey questions are described below, in Table S4. All P*Am* derived data fields are described in Table S5. The original and derived data fields from the ABF survey are described in Table S6.

#### Project *Apis m.* Survey Binning

To accompany Tables S4 and S5, we include the following narrative descriptions of the process through which open-ended responses were cleaned and categorized into derived data fields.

*Predicted Causes of Colony Losses, derived from "Predicted_Cause"*

**Survey question:** Recognizing this is just a guess, what factors caused your colony losses this year? ["Predicted_Cause"]

**Possible responses:** Pesticides, *Varroa* mites, Queen failure, Starvation, Weather, Disease, Other (open category); A single respondent was counted in multiple categories if more than one choice was made. Responses with a version of "all of the above" had each category selected.

**Binning:** The final categories in the analysis were: Other, Pesticides, *Varroa* mites, Queen failure, Food, Weather, Disease, and Unknown. Responses were binned into one of the final categories where appropriate, as captured in the binary variables beginning with "cause_" (see Table S5):

- ***Varroa***: Mites or *Varroa.*
- **Pesticides**: Pesticides or chemical applications other than *Varroa* treatments.
- **Pathogens**: Disease, European foulbrood (EFB), Nosema, and/or viruses.
- **Queen Failure**: Queen failure, loss, issues, problems, or longevity.
- **Weather**: Heat, drought, hurricanes, floods, or moisture (other than hive moisture).
- **Food**: Nutrition and starvation.
- **Unknown**: Beekeeper indicated that they did not know the cause of the loss, colony collapse disorder, or colony absconded.
- **Other**: Factors that did not fit into the other categories.

*Varroa destructor mite treatments, derived from "Mite_Treatment_Freq"*

**Survey question:** How often did you treat for mites and what treatment was used from June-December, 2024? ["Mite_Treatment_Freq"]

**Possible responses:** Open-ended.

**Binning:** responses were binned into binary variables beginning with "trt_" (see Table S5), and then combined into the below categories for analysis.

- **Amitraz Only:** Includes all entries where a product was used where the active ingredient was amitraz and no other product use was specified.

- **Product Unknown:** The frequency of a treatment was provided, but not the type of product used. Respondents may have indicated an unlisted product and another listed product.
- **Amitraz+:** Respondent indicated they used an amitraz-based product and at least one other chemistry known to be effective against *Varroa* mites (a "Non-Amitraz Known" product).
- **Non-Amitraz Known**: Respondent listed only treatment(s) that were *not* amitraz-based. Includes products with one of the following active ingredients: oxalic acid, formic acid, or thymol.
- **None**: Respondent indicated that they did not apply a chemical treatment within the June to December 2024 timeframe. Respondents may have used management to control mites, including *Varroa* resistant stock, splitting, brood breaks, drone brood removal, or heat.

*Supplemental feeding of protein, derived from "Supplemental_Feed_Protein"*

**Survey question:** Did you supplementally feed your bees protein- pollen, substitutes, etc.? If yes, what and when? ["Supplemental_Feed_Protein"]

**Possible responses:** The responses were open-ended. Example responses included "Yes" or "No" for supplemental protein feeding. For responses to the question "If yes, what and when", some respondents described the detailed protein feeding regimen ("Yes. In the cold weather dropped in protein patty and I provide pollen substitute in all my B yards") while others only mentioned the months ("Yes, Dec & Jan"). Type of protein feeding responses varied from "pollen patties" to "pollen substitute" to detailed product and feeding descriptions or none at all.

**Binning:** responses were binned into binary variables beginning with "Protein_" (see Table S5)

- **Protein Feed:** "Yes" or "No" feeding responses were directly included. For descriptive answers, any explanation of protein feeding type or frequency was considered a "Yes".
- **Protein Frequency:** The number of times supplemental proteins were fed to the honey bee colonies, as recorded by the survey respondents, has been grouped into four frequencies. For those respondents who did not feed proteins (pollen or substitutes), the responses were grouped as "0". For just one supplemental protein feeding between June 2024 - March 2025 (the timeline for survey respondents), they were grouped as "1". Two supplemental protein feedings in any form are grouped as "2" and three or

more than three protein supplemental feedings have been grouped as “3+”. When survey respondents described the months and number of feedings, the data was directly recorded. When only seasons or months were included, the responses were grouped according to the seasons. The seasons considered in the survey included Spring (April - June), Summer (July - September), Fall (October - December) and Winter (January - March).

- **Type:** The type of protein supplementation given to the colonies based on the survey responses. The responses varied between commercial diets or homemade patties or mixed supplementations. Anything that was supplemented and was not a protein is coded as "no" for protein supplementation type.
- **Season:** Spring (April -June), Summer (July - September), Fall (October - December) and Winter (January - March).
- **Company name:** If the respondents indicated names of any commercial protein supplements used, that has been binned as well.
- **Amount:** Weights in pounds as per the respondents.
- **Probiotics:** Any commercial probiotic used by the respondent and described in the open-ended answer write-ins.
- **ProComp:** Any probiotic company names the respondents provided.
- **Supplements:** Any non-protein supplements and non-probiotic supplements used.

*Supplemental feeding of carbohydrates, derived from “Supplemental_Feed_Sugar”*

**Survey question:** Did you supplementally feed your bees sugar or syrup? If yes, what months? [“Supplemental_Feed_Sugar”]

**Possible responses:** The responses were open-ended. Example responses included “yes or no” for feeding. For responses to the question “If yes, what months”, some respondents described the detailed sugar or syrup feeding regimen ("Yes I fed syrup in the fall a couple of times”) while some others only mentioned the months (“June, Aug, Sept, Oct, Dec, Jan”). Type of sugar feeding responses varied from “sugar” to “syrup” to detailed product descriptions or none at all.

**Binning:** responses were binned into binary variables beginning with “Sugar_” (see Table S5)

- **Sugar Feed:** “Yes” or “No” feeding responses were directly included. For descriptive answers, any explanation of sugar feeding type or frequency was considered a “Yes”.
- **Sugar Frequency:** The number of times supplemental carbohydrates were fed to the honey bee colonies, as recorded by the survey respondents, has been grouped into four frequencies. Less than four supplemental feedings of sugar in any form are grouped as “<4” and more than four carbohydrate supplemental feedings have been grouped as “>4”. When survey respondents described the months and number of feedings, the data was directly recorded. When only seasons or months have been included, the responses were grouped according to the seasons. The seasons considered in the survey included Spring (April - June), Summer (July - September), Fall (October - December) and Winter (January - March).
- **Company name:** If survey respondents mentioned any commercial sugar supplement by name, that has been included.
- **Amount:** Quantity in gallons of sugar supplement fed.
- **Probiotics:** Any commercial probiotic used by the respondent and described in the open-ended answer write-ins.
- **ProComp:** Any probiotic company names the respondents provided.
- **Supplements:** Any non-carbohydrate and non-probiotics commercial supplements used.
- **Ratio:** Ratio of sugar to water or different blends used and as reported.
- **Overwintering:** Respondent mention of overwintering indoors or outdoors.

**Supplemental Tables**

Table S1. Significant results for models investigating the quality of covariates and loss data. Independent variables include mean minimum winter temperature (November-February), total active season precipitation (in., June-September), and mean daily active season temperature (June-September). Loss variables investigated include Winter, Summer, and Net Losses.

| Model | df | $\chi^2$ | Pr(>$\chi^2$) |
|---|---|---|---|
| **Winter Loss v Min Winter Temp** | | | |
| Commercial | 1 | 2.24 | 0.135 |
| Sideliner | 1 | 5.273 | **0.022** |
| Hobbyist | 1 | 12.446 | **4.19E-04** |
| **Net Loss v Total Precipitation** | | | |
| Commercial | 1 | 0.372 | 0.542 |
| Sideliner | 1 | 7.315 | **0.007** |
| Hobbyist | 1 | 7.636 | **0.006** |
| **Net Loss v Active Season Temp** | | | |
| Commercial | 1 | 2.487 | 0.115 |
| Sideliner | 1 | 6.354 | **0.012** |
| Hobbyist | 1 | 10.136 | **0.001** |
| **Winter Loss v Summer Loss** | | | |
| Commercial | 1 | 15.795 | **7.06E-05** |
| Sideliner | 1 | 4.769 | **0.029** |
| Hobbyist | 1 | 9.883 | **0.002** |

Table S2. Self-reported miticide products used from June to December 2024 in the P*Am* survey. Beekeepers often referred to products by the active ingredient and not by a product name. Totals may differ from Figure 2b as these data include all the beekeepers responding to the question about *Varroa* treatment rather than the subset for the paired analyses. Of the beekeepers that reported a specific method of oxalic acid use, 27 used the dribble method, 55 used extended release (including VarroxSan), and 145 used the vaporizing/fumigating method.

| | **n** | **Amitraz** | **Formic Acid** | **Oxalic Acid** | **Thymol** | **Other Products *** | **Product not provided** | **No chemical treatment** |
|---|---|---|---|---|---|---|---|---|
| Commercial (over 500 hives) | 275 | 133 | 73 | 115 | 45 | 8 | 88 | 3 |
| Sideliner (50-500 hives) | 160 | 43 | 40 | 85 | 36 | 4 | 43 | 9 |
| Hobbyist (1-49 hives) | 383 | 51 | 73 | 136 | 42 | 8 | 109 | 79 |
| NA | 3 | 0 | 0 | 0 | 0 | 0 | 1 | 2 |
| Total | 821 | 227 | 186 | 336 | 123 | 20 | 241 | 93 |

* Hopguard, coumaphos, and fluvalinate

Table S3. Self-reported frequency of miticide usage from June through December 2024 in the P*Am* survey.

| Beekeeper Class | n | Mean frequency of treatments | Standard deviation | Range |
|---|---|---|---|---|
| Commercial (over 500 hives) | 236 | 4.67 | 3.11 | 0 to 22 |
| Sideliner (50-500 hives) | 147 | 3.53 | 2.44 | 0 to 16 |
| Hobbyist (1-49 hives) | 345 | 2.12 | 2.37 | 0 to 21 |
| NA | 3 | 7.00 | 12.10 | 0 to 21 |

Table S4. Project *Apis m.* Original Survey Fields.

| Field | Description |
|---|---|
| **ID** | A unique numeric identifier for each survey response |
| **Name** | [Redacted] Your name and operation (optional) |
| **Submission_Type** | [Redacted] Is this the first submission of your data using this form? If this is an update, please share info for us to identify your last entry. (name, and what changes occurred) |
| **Loss_Summer_Fall** | [Redacted] What is your estimated colony loss % from June 2024 to winter? [Note, redacted because some responses contain personally identifiable information. See Loss_Summer_Fall_Clean.] |
| **Loss_Winter** | [Redacted] What is your estimated colony loss % during winter? [See Loss_Winter_Clean.] |
| **Loss_Post_Emergence** | [Redacted] What is your estimated colony loss % from first check post-winter to two months later? [See Loss_Post_Emergence_Clean] |
| **Colony_Relative_Size** | How would you categorize your overwintered colony size this year? |
| **Location_Summer** | [Redacted] Where were your bees June-Fall 2024? |
| **Location_Winter** | [Redacted] Where were your bees over winter, 2024? |
| **Location_Rebuild** | [Redacted] Where were your bees from emergence until Feb 1? |
| **Percent_Almonds** | What percentage of your bees in early February went to pollinate almonds this year? |
| **Predicted_Cause** | Recognizing this is just a guess, what factors caused your colony losses this year? |
| **Pesticide_Details** | If you suspect pesticide exposure contributed to your losses, please describe any details. (timing, region, crop, application of products used, etc) |
| **Mite_Treatment_Freq** | [Redacted] How often did you treat for mites and with what from June-December, 2024? |
| **Mite_Count** | What were your average Varroa mite levels in the Fall? |
| **Other_Pests** | Did you encounter other pests (e.g., small hive beetles, wax moths)? |
| **Supplemental_Feed_Sugar** | Did you supplementally feed your bees sugar or syrup? if yes, what months? |
| **Supplemental_Feed_Protein** | Did you supplementally feed your bees protein- pollen, substitutes, etc? If yes, what and when? |
| **Queen_Replacement** | What percentage of your queens were replaced due to queen health from June-winter 2024? |
| **Percent_Indoor** | What percentage of your bees were overwintered indoors versus outdoors? |
| **Percent_Loss_Indoor_Outdoor** | If you had some indoor and outdoor, what were the % losses? |
| **Beekeeper_Class** | What is the size of your business? |
| **Financial_Concern** | If your average business financial concern is a 5, rank your business financial concerns this year. |
| **Contact_OK** | [Redacted] Additional information could become helpful. Will you allow P*Am* or researchers to contact you and gather follow up information? Please reply with "yes" or "no", your contact information, and the b… |

Table S5. Derived Data Fields.

| Field | Description |
|---|---|
| **Loss_Summer_Fall_Clean** | Cleaned version of Loss_Summer_Fall, stored as text |
| **Loss_Summer_Fall_n** | Cleaned version of Loss_Summer_Fall, converted to numerical value between 0-1 (i.e., proportional loss) |
| **Loss_Winter_Clean** | Cleaned version of Loss_Winter, stored as text |
| **Loss_Winter_n** | Cleaned version of Loss_Winter, converted to numerical value between 0-1 (i.e., proportional loss) |
| **Loss_Post_Emergence_Clean** | Cleaned version of Loss_Post_Emergence, stored as text |
| **Loss_Post_Emergence_n** | Cleaned version of Loss_Post_Emegence, converted to numerical value between 0-1 (i.e., proportional loss) |
| **Colony_Relative_Size_Clean** | Cleaned version of "Colony_Relative_Size" consisting of the following values: |
| **Updated** | Whether any of the percent loss fields have been updated during cleaning (values are either "original" or "updated") |
| **Loss_Summer_Fall_Updated** | Whether Loss_Summer_Fall was updated during cleaning (TRUE [i.e., was updated] or FALSE [i.e., was not updated]) |
| **Loss_Winter_Updated** | Whether Loss_Winter was updated during cleaning (TRUE [i.e., was updated] or FALSE [i.e., was not updated]) |
| **Loss_Post_Emergence_Updated** | Whether Loss_Post_Emergence was updated during cleaning (TRUE [i.e., was updated] or FALSE [i.e., was not updated]) |
| **Duplicate** | Whether the row is identified as a duplicate entry to be removed during analysis ("keep" or "remove") |
| **Summer_Climate_Zone_1** | US Climate Zone corresponding to the primary, or first listed, state location for Summer 2024, extracted from "Location_Summer." Climate Zone classifications developed by NOAA's National Centers for Environmental Information (Karl and Koss, 1984). See here: https://www.ncei.noaa.gov/access/monitoring/reference-maps/us-climate-regions |
| **Winter_Climate_Zone_1** | US Climate Zone corresponding to the primary, or first listed, state location for Winter 2024, extracted from "Location_Winter." See Climate Zone citation for “Summer_Climate_Zone_1”. |
| **Rebuild_Climate_Zone_1** | US Climate Zone corresponding to the primary, or first listed, state location for Rebuild 2025, extracted from "Location_Rebuild." See Climate Zone citation for “Summer_Climate_Zone_1”. |
| **Net_Survival** | The proportion of all colonies that survived through all three time periods. Calculated as follows: = (1 - Loss_Summer_Fall_n) * (1 - Loss_Winter_n) * (1 - Loss_Post_Emergence_n) |
| **Net_Loss** | The overall loss rate through all three time periods. Calculated as follows: = (1- Net_Survival) |

Table S5, continued. Derived Data Fields.

| Field | Description |
|---|---|
| ***Supplemental feeding:*** | |
| **Protein_Feed** | Yes or No feeding |
| **Protein_Type** | Commercial diet or homemade patty or mixed supplementation for protein patties for example. Anything that is not a protein is coded as "no" for protein supplementation. |
| **Protein_Probiotics** | Whether commercial probiotic has been fed or not and how many times even if the probiotic has been fed with or without other supplementation |
| **Protein_Frequency** | How many times in the year or how many months indicated in survey for protein supplements |
| **Protein_Spring** | Protein supplement frequency in Spring (April -June) |
| **Protein_Summer** | Protein supplement frequency in Summer (July - September) |
| **Protein_Fall** | Protein supplement frequency in Fall (October - December) |
| **Protein_Winter** | Protein supplement frequency in Winter (January - March) |
| **Protein_Amount** | Weights in pounds as per the survey results |
| **Protein_Company** | As survey indicated for protein patties |
| **Protein_ProComp** | Probiotic company name as the survey reports suggested |
| **Protein_Supplements** | Any non-protein supplements and non-probiotic supplements used |
| **Protein_Notes** | Anything else the survey has |
| **Sugar_Feed** | Yes or No feeding |
| **Sugar_Type** | Commercial formulation or homemade formulation or mixed formulations. Anything that is not a sugar supplement is coded as "no" for these tables and also sugar water or sugar syrup are both called sugar water |
| **Sugar_Probiotics** | Whether commercial probiotic has been fed or not and how many times even if the probiotic has been fed with or without other supplementation |
| **Sugar_Frequency** | How many times in the year or how many months indicated in survey for sugar supplements |
| **Sugar_Spring** | Sugar supplement frequency in Spring (April -June) |
| **Sugar_Summer** | Sugar supplement frequency in Summer (July - September) |
| **Sugar_Fall** | Sugar supplement frequency in Fall (October - December) |
| **Sugar_Winter** | Sugar supplement frequency in Winter (January - March) |
| **Sugar_Amount** | Weights in gallons as per the survey results |
| **Sugar_Ratio** | Ratio of sugar to water or different blends used and as reported in the beekeeper surveys |
| **Sugar_Company** | Company name as survey indicated for sugar supplements |
| **Sugar_ProComp** | Probiotic company name as the survey reports suggested |
| **Sugar_Supplements** | Any non-carbohydrate and non-probiotics commercial supplements used |
| **Sugar_Notes** | Any other relevant notes in the survey |
| **Indoor_overwintering** | If beekeeper used indoor storage facilities as indicated in the survey |

Table S5, continued. Derived Data Fields

| Field | Description |
|---|---|
| ***Varroa destructor* mite treatments** | |
| **trt_category** | [Primary] Method of varroa management the beekeeper reported using: None (treatment free), Treatment Unknown (did not state or was confusing; most are probably amitraz), Product Unknown (Hoperthermia, mitral, tramisol & mustard), Management (brood breaks, varroa resistant stocks, splits, and/or heat), and/or a Treatment (chemical treatment stated). Note: this and all fields starting with "trt_" are derived from the original survey question captured in "Mite_Treatment_Freq" above. |
| **trt_essoils** | Beekeeper reported using "essential oils" |
| **trt_apivar** | Beekeeper used Apivar |
| **trt_amiflex** | Beekeeper used AmiFlex |
| **trt_amitraz** | Beekeeper used "amitraz" and did not specify Apivar or AmiFlex |
| **trt_amitraz_all** | [Primary] Beekeeper used amitraz (Apivar, AmiFlex, and/or amitraz) |
| **trt_oxalic** | Beekeeper used oxalic acid and did not specify a specific product or application |
| **trt_varroxsan** | Beekeeper used VarroxSan |
| **trt_oxalic_extendedrelease** | Beekeeper used extended-release oxalic acid and did not identify the product as VarroxSan |
| **trt_oxalic_vapor** | Beekeeper used oxalic acid vaporization/fumigation |
| **trt_oxalic_dribble** | Beekeeper used oxalic acid dribble |
| **trt_oxalic_all** | [Primary] Beekeeper used any form of oxalic acid |
| **trt_formicmaqs** | Beekeeper used Mite Away Quick Strips |
| **trt_formicpro** | Beekeeper used Formic Pro |
| **trt_formic** | Beekeeper used formic acid and did not identify a specific product |
| **trt_formic_all** | [Primary] Beekeeper used any form of formic acid |
| **trt_apiguard** | Beekeeper used Apiguard |
| **trt_apilifevar** | Beekeeper used Api LifeVar |
| **trt_thymol** | Beekeeper used thymol and did not identify a specific product |
| **trt_thymol_all** | [Primary] Beekeeper used any form of thymol |
| **trt_hopguard** | Beekeeper used HopGuard |
| **trt_fluvalinate** | Beekeeper used fluvalinate/Apistan |
| **trt_coumaphos** | Beekeeper used coumaphos/ CheckMite+ |
| **trt_product_not_listed** | [Primary] Beekeeper did not specify the type(s) of miticide used to treat varroa, but did indicate frequency of product usage. |
| **trt_treatment_type** | Binned response of trt_amitraz_all, trt_product_not_listed, trt_oxalic_all, trt_formic_all, trt_thymol_all, trt_none_management. See Supplemental Methods under *Varroa destructor mite treatments* for list and explanation of binned responses. |

Table S5, continued. Derived Data Fields

| Field | Description |
|---|---|
| **trt_none_management** | [Primary] - Beekeeper did not apply a chemical treatment from June to December 2024. Some respondents used management to control mites (varroa resistant stock, splitting, brood breaks, drone brood removal, or heat). |
| **trt_frequency_not_listed** | Varroa treatment product(s) included, but not the time of year they were used nor the use frequency; TRUE or NA |
| **trt_freq_approximate** | Frequency of treatment application approximated based on pre/post honey flow or season (spring, summer, fall) or the number of products listed; TRUE or NA |
| **trt_freq_full_year** | Estimated frequency of mite treatment use for beekeepers who reported miticide data for a full year instead of June to Dec; if data were incomplete and the beekeeper did not specify the timing for all treatments, then no treatments were included in the four seasons |
| **trt_freq_JunDec** | Number of times a beekeeper treated for varroa from June to December 2024; partial numbers were used if a beekeeper gave a range and the average of that range was not a whole number (e.g., 3-4 was entered as 3.5), if the 12-month number was reduced, or if they did not treat all their bees |
| **trt_freq_spring_Jun.** | Number of times a beekeeper reported treating for varroa in June 2024 and the product used if available; only June was included because April and May were not part of the survey question |
| **trt_freq_summer_Jul_Sep.** | Number of times a beekeeper reported treating for varroa from July to September 2024 and the treatment used if available |
| **trt_freq_fall_Oct_Dec** | Number of times a beekeeper reported treating for varroa from October to December 2024 and the treatment used if available |
| **trt_freq_amitraz** | Number of times a beekeeper reported using amitraz |
| **trt_freq_apivar** | Number of times a beekeeper reported using Apivar |
| **trt_freq_amiflex** | Number of times a beekeeper reported using AmiFlex |
| **trt_freq_oa** | Number of times a beekeeper reported using oxalic acid (not specifying vapor, dribble, VarroxSan, or extended release) |
| **trt_freq_oav** | Number of times a beekeeper reported using oxalic acid vaporization/fumigation |
| **trt_freq_oae** | Number of times a beekeeper reported using oxalic acid extended release (not specifying VarroxSan) |
| **trt_freq_oadribble** | Number of times a beekeeper reported using oxalic acid dribble |
| **trt_freq_varroxsan** | Number of times a beekeeper reported using VarroxSan |
| **trt_freq_apiguard** | Number of times a beekeeper reported using Apiguard |
| **trt_freq_thymol** | Number of times a beekeeper reported using an unspecified thymol product |
| **trt_freq_formicpro** | Number of times a beekeeper reported using Formic Pro |

Table S5, continued. Derived Data Fields

| Field | Description |
|---|---|
| **trt_freq_formic** | Number of times a beekeeper reported using an unspecified formic acid product |
| **trt_freq_maqs** | Number of times a beekeeper reported using Mite Away Quick Strips |
| **trt_freq_apivlifevar** | Number of times a beekeeper reported using ApiLife Var |
| **trt_freq_ess_oils** | Number of times a beekeeper reported using essential oils |
| **trt_freq_fluvalinate** | Number of times a beekeeper reported using Apistan or fluvalinate |
| **trt_freq_hopguard** | Number of times a beekeeper reported using Hopguard |
| **trt_freq_checkmite** | Number of times a beekeeper reported using CheckMite+ or coumaphos |
| **N_Treatments** | Number of mite treatments, drawn from " Mite_Treatment_Freq", containing responses to the question "How often did you treat for mites and with what from June-December, 2024". |
| **TypeMiteTreat** | Code mite treatment data, drawn from " Mite_Treatment_Freq", containing responses to the question "How often did you treat for mites and with what from June-December, 2024". Values include: "AM", "AMFO", "AMFOTH", "AMOT", "AMOX", "AMOXFO", "AMOXFOHG", "AMOXFOTH", "AMOXHG", "AMOXTH", "AMOXTHOT", "AMTH", "AMTHFL", "AXOXTH", "FL", "FO", "FOFL", "FOHG", "FOTH", "FOTHHG", "HG", "MU", "NO", "OA", "OAHG", "OAOT", "OT", "OX", "OXFO", "OXFOHG", "OXFOTH", "OXTH", "TH", "THHG", "UN" |
| **Amitraz_Use** | Categorical variable indicating whether amitraz was used for mite treatment. Values include: "Likely", "No", or "Yes". Drawn from "Mite_Treatment_Freq", containing responses to the question "How often did you treat for mites and with what from June-December, 2024". |
| **Mite_Levels** | Categorical, binned data drawn from "Mite_Count", which contains responses to the question, "What were your average Varroa mite levels in the Fall?". Values include: "0-1m", "2-5m", "5+m", "NoTest" |
| **Pest_Detected** | Whether responding beekeepers detected another pest, drawn from "Other_Pest", responding to question, "Did you encounter other pests (e.g., small hive beetles, wax moths)?" Values include: "No", "Yes", or "NA" |
| **Which_Pest** | Cleaned data indicating which other pests besides Varroa mites (if any) were detected by responding beekeepers. Drawn from "Other_Pest", responding to question, "Did you encounter other pests (e.g., small hive beetles, wax moths)? Values include: "NA", "OT", "SHB", "SHBOT", "SHBWM", "SHBWMOT", "UN", "WM", "WMOT". |
| **queenloss** | Binned data on queen loss provided in "Queen_Replacement", based on the question "What percentage of your queens were replaced due to queen health from June-winter 2024?". Values are: "None", "<10%", "10-30%", "30-50%", or ">50%". |
| **WinterMode** | Categorial data on where bees spent winter, drawn from "Percent_Indoor", based on the survey question "What percentage of your bees were overwintered indoors versus outdoors?" Values are "Outside", "Mixed", or "Shed". |

Table S5, continued. Derived Data Fields

| Field | Description |
|---|---|
| ***Predicted colony loss causes*** | |
| **NetCause** | A combined field summarizing the fields that start with "cause_" and are derived from "Predicted_Cause." The values are 4-letter codes corresponding to the specific causes: "DISE" (i.e., disease or virus), "FOOD" (i.e., starvation or nutrition), "PCID" (i.e., pesticides), "PEST" (i.e., pests), "QUEE" (i.e., queen failure), "VARR" (i.e., varroa), "WEAT" (i.e., weather or hurricane). Does not include the following causes: "unknown", "ccd", "colony_size", "abscond", "no_losses", or "other". |
| **NetCauseRED** | Same as NetCause, but listed as "MULT" when 3 or more causes are mentioned. |
| **cause_unknown** | Cause of loss unknown to beekeeper; includes colony collapse disorder (CCD), absconding, and various iterations of "don't know"; entry in file: T |
| **cause_pesticides** | [Primary] Beekeeper indicated pesticide exposure was a possible cause of losses; does not include beekeeper-applied miticides; one of the options in the P*Am* survey; entry in file: T |
| **cause_varroa** | [Primary] Beekeeper indicated varroa mites were a possible cause of losses; one of the options in the P*Am* survey; entry in file: T |
| **cause_queen_failure** | [Primary] Beekeeper indicated queen failure was a possible cause of losses, including any mention of failure, issues, problems, or longevity specific to the queen; one of the options in the P*Am* survey; entry in file: T |
| **cause_food** | [Primary] Beekeeper indicated either starvation or poor nutrition as possible causes of losses; entry in file: T |
| **cause_weather** | [Primary] Beekeeper indicated weather was a possible cause of losses; includes heat, drought, hurricanes, floods, or moisture (other than hive moisture); one of the options in the P*Am* survey; entry in file: T |
| **cause_pathogens** | [Primary] Beekeeper indicated disease, European foulbrood (EFB), nosema, and/or viruses (viruses may or may not be associated with varroa, n = 27) as a possible cause of loss; "Disease" was one of the options presented in the P*Am* survey; entry in file: T |
| **cause_no_losses** | [Primary] Beekeeper indicated that they had no losses during the time period and therefore did not have a cause of loss; entry in file: T |
| **cause_other** | [Primary] Beekeeper indicated that another factor was a possible cause of loss; included dwindling, pests, parasites, colony size, labor/costs , moisture, tracheal mites, chemtrails, electromagnetic fields, sinkhole, genetics, robbing, equipment failure, Geoengineering, too much food, air quality, a fog, failure of endemic species, absconding/death due to varroa treatment, beekeeper error/care, relocated hive, and swarming; entry in file: T |

Table S5, continued. Derived Data Fields

| Field | Description |
|---|---|
| ***Weather information (precipitation and temperature)*** | |
| **precip_mean_Jan** | Mean daily precipitation for the month of January, for the state listed in 'Summer_State_1'. Data provided by the PRISM Climate Group, Oregon State University. https://www.prism.oregonstate.edu/ |
| **precip_mean_Feb** | Mean daily precipitation for the month of February, for the state listed in 'Summer_State_1'. Data provided by the PRISM Climate Group, Oregon State University. https://www.prism.oregonstate.edu/ |
| **precip_mean_June** | Mean daily precipitation for the month of June, for the state listed in 'Summer_State_1'. Data provided by the PRISM Climate Group, Oregon State University. https://www.prism.oregonstate.edu/ |
| **precip_mean_July** | Mean daily precipitation for the month of July, for the state listed in 'Summer_State_1'. Data provided by the PRISM Climate Group, Oregon State University. https://www.prism.oregonstate.edu/ |
| **precip_mean_Aug** | Mean daily precipitation for the month of August, for the state listed in 'Summer_State_1'. Data provided by the PRISM Climate Group, Oregon State University. https://www.prism.oregonstate.edu/ |
| **precip_mean_Sep** | Mean daily precipitation for the month of September, for the state listed in 'Summer_State_1'. Data provided by the PRISM Climate Group, Oregon State University. https://www.prism.oregonstate.edu/ |
| **precip_mean_Oct** | Mean daily precipitation for the month of October, for the state listed in 'Summer_State_1'. Data provided by the PRISM Climate Group, Oregon State University. https://www.prism.oregonstate.edu/ |
| **precip_mean_Nov** | Mean daily precipitation for the month of November, for the state listed in 'Summer_State_1'. Data provided by the PRISM Climate Group, Oregon State University. https://www.prism.oregonstate.edu/ |
| **precip_mean_Dec** | Mean daily precipitation for the month of December, for the state listed in 'Summer_State_1'. Data provided by the PRISM Climate Group, Oregon State University. https://www.prism.oregonstate.edu/ |
| **precip_sum_Jan** | Total precipitation for the month of January, for the state listed in 'Summer_State_1'. Data provided by the PRISM Climate Group, Oregon State University. https://www.prism.oregonstate.edu/ |
| **precip_sum_Feb** | Total precipitation for the month of February, for the state listed in 'Summer_State_1'. Data provided by the PRISM Climate Group, Oregon State University. https://www.prism.oregonstate.edu/ |
| **precip_sum_June** | Total precipitation for the month of June, for the state listed in 'Summer_State_1'. Data provided by the PRISM Climate Group, Oregon State University. https://www.prism.oregonstate.edu/ |
| **precip_sum_July** | Total precipitation for the month of July, for the state listed in 'Summer_State_1'. Data provided by the PRISM Climate Group, Oregon State University. https://www.prism.oregonstate.edu/ |

Table S5, continued. Derived Data Fields

| Field | Description |
|---|---|
| **precip_sum_Aug** | Total precipitation for the month of August, for the state listed in 'Summer_State_1'. Data provided by the PRISM Climate Group, Oregon State University. https://www.prism.oregonstate.edu/ |
| **precip_sum_Sep** | Total precipitation for the month of September, for the state listed in 'Summer_State_1'. Data provided by the PRISM Climate Group, Oregon State University. https://www.prism.oregonstate.edu/ |
| **precip_sum_Oct** | Total precipitation for the month of October, for the state listed in 'Summer_State_1'. Data provided by the PRISM Climate Group, Oregon State University. https://www.prism.oregonstate.edu/ |
| **precip_sum_Nov** | Total precipitation for the month of November, for the state listed in 'Summer_State_1'. Data provided by the PRISM Climate Group, Oregon State University. https://www.prism.oregonstate.edu/ |
| **precip_sum_Dec** | Total precipitation for the month of December, for the state listed in 'Summer_State_1'. Data provided by the PRISM Climate Group, Oregon State University. https://www.prism.oregonstate.edu/ |
| **tmean_Jan** | Mean daily temperature for the month of January, for the state listed in 'Summer_State_1'. Data provided by the PRISM Climate Group, Oregon State University. https://www.prism.oregonstate.edu/ |
| **tmean_Feb** | Mean daily temperature for the month of February, for the state listed in 'Summer_State_1'. Data provided by the PRISM Climate Group, Oregon State University. https://www.prism.oregonstate.edu/ |
| **tmean_June** | Mean daily temperature for the month of June, for the state listed in 'Summer_State_1'. Data provided by the PRISM Climate Group, Oregon State University. https://www.prism.oregonstate.edu/ |
| **tmean_July** | Mean daily temperature for the month of July, for the state listed in 'Summer_State_1'. Data provided by the PRISM Climate Group, Oregon State University. https://www.prism.oregonstate.edu/ |
| **tmean_Aug** | Mean daily temperature for the month of August, for the state listed in 'Summer_State_1'. Data provided by the PRISM Climate Group, Oregon State University. https://www.prism.oregonstate.edu/ |
| **tmean_Sep** | Mean daily temperature for the month of September, for the state listed in 'Summer_State_1'. Data provided by the PRISM Climate Group, Oregon State University. https://www.prism.oregonstate.edu/ |
| **tmean_Oct** | Mean daily temperature for the month of October, for the state listed in 'Summer_State_1'. Data provided by the PRISM Climate Group, Oregon State University. https://www.prism.oregonstate.edu/ |
| **tmean_Nov** | Mean daily temperature for the month of November, for the state listed in 'Summer_State_1'. Data provided by the PRISM Climate Group, Oregon State University. https://www.prism.oregonstate.edu/ |

Table S5, continued. Derived Data Fields

| Field | Description |
| --- | --- |
| **tmean_Dec** | Mean daily temperature for the month of December, for the state listed in 'Summer_State_1'. Data provided by the PRISM Climate Group, Oregon State University. https://www.prism.oregonstate.edu/ |
| **tmin_Jan** | Mean daily minimum temperature for the month of January, for the state listed in 'Summer_State_1'. Data provided by the PRISM Climate Group, Oregon State University. https://www.prism.oregonstate.edu/ |
| **tmin_Feb** | Mean daily minimum temperature for the month of February, for the state listed in 'Summer_State_1'. Data provided by the PRISM Climate Group, Oregon State University. https://www.prism.oregonstate.edu/ |
| **tmin_June** | Mean daily minimum temperature for the month of June, for the state listed in 'Summer_State_1'. Data provided by the PRISM Climate Group, Oregon State University. https://www.prism.oregonstate.edu/ |
| **tmin_July** | Mean daily minimum temperature for the month of July, for the state listed in 'Summer_State_1'. Data provided by the PRISM Climate Group, Oregon State University. https://www.prism.oregonstate.edu/ |
| **tmin_Aug** | Mean daily minimum temperature for the month of August, for the state listed in 'Summer_State_1'. Data provided by the PRISM Climate Group, Oregon State University. https://www.prism.oregonstate.edu/ |
| **tmin_Sep** | Mean daily minimum temperature for the month of September, for the state listed in 'Summer_State_1'. Data provided by the PRISM Climate Group, Oregon State University. https://www.prism.oregonstate.edu/ |
| **tmin_Oct** | Mean daily minimum temperature for the month of October, for the state listed in 'Summer_State_1'. Data provided by the PRISM Climate Group, Oregon State University. https://www.prism.oregonstate.edu/ |
| **tmin_Nov** | Mean daily minimum temperature for the month of November, for the state listed in 'Summer_State_1'. Data provided by the PRISM Climate Group, Oregon State University. https://www.prism.oregonstate.edu/ |
| **tmin_Dec** | Mean daily minimum temperature for the month of December, for the state listed in 'Summer_State_1'. Data provided by the PRISM Climate Group, Oregon State University. https://www.prism.oregonstate.edu/ |
| **tmax_Jan** | Mean daily maximum temperature for the month of January, for the state listed in 'Summer_State_1'. Data provided by the PRISM Climate Group, Oregon State University. https://www.prism.oregonstate.edu/ |
| **tmax_Feb** | Mean daily maximum temperature for the month of February, for the state listed in 'Summer_State_1'. Data provided by the PRISM Climate Group, Oregon State University. https://www.prism.oregonstate.edu/ |
| **tmax_June** | Mean daily maximum temperature for the month of June, for the state listed in 'Summer_State_1'. Data provided by the PRISM Climate Group, Oregon State University. https://www.prism.oregonstate.edu/ |

Table S5, continued. Derived Data Fields

| Field | Description |
|---|---|
| **tmax_July** | Mean daily maximum temperature for the month of July, for the state listed in 'Summer_State_1'. Data provided by the PRISM Climate Group, Oregon State University. https://www.prism.oregonstate.edu/ |
| **tmax_Aug** | Mean daily maximum temperature for the month of August, for the state listed in 'Summer_State_1'. Data provided by the PRISM Climate Group, Oregon State University. https://www.prism.oregonstate.edu/ |
| **tmax_Sep** | Mean daily maximum temperature for the month of September, for the state listed in 'Summer_State_1'. Data provided by the PRISM Climate Group, Oregon State University. https://www.prism.oregonstate.edu/ |
| **tmax_Oct** | Mean daily maximum temperature for the month of October, for the state listed in 'Summer_State_1'. Data provided by the PRISM Climate Group, Oregon State University. https://www.prism.oregonstate.edu/ |
| **tmax_Nov** | Mean daily maximum temperature for the month of November, for the state listed in 'Summer_State_1'. Data provided by the PRISM Climate Group, Oregon State University. https://www.prism.oregonstate.edu/ |
| **tmax_Dec** | Mean daily maximum temperature for the month of December, for the state listed in 'Summer_State_1'. Data provided by the PRISM Climate Group, Oregon State University. https://www.prism.oregonstate.edu/ |

Table S6. American Beekeeping Federation (ABF) Survey Fields.

| Field | Description |
|---|---|
| **ID** | A unique numeric identifier for each survey response |
| **N_Colonies_Dec_2024** | Original survey question: "How many colonies did you have December 1, 2024". Categorical variable with the following options: "Less than 500"; "Between 500 and 1,000"; "Between 1,000 and 3,000"; "Between 3,000 and 10,000"; or "More than 10,000" |
| **Hive_Size** | Original survey question: "What size are you hives?" Categorical variable with the following options: "Single Deep, 8 Frame"; "Double Deep, 8 Frame"; "Single Deep, 10 Frame"; or "Double Deep, 10 Frame" |
| **N_Colonies_Lost_since_Dec_2024** | Original survey question: "How many colonies have you lost since December 1, 2024?" Open-ended response. |
| **Location_June_2024** | Original survey question: "Where were your hives June 2024?" |
| **Overwinter_Method** | Original survey question: "How were your hives overwintered?" Categorical variable with the following options: "Shed"; "Outside" |
| **Mite_Treatment_Details** | Original survey question: "What is your routine mite treatment? # and kind of treatment" Open-ended response. |
| **Mite_Treatment_Efficacy** | Original survey question: "Was your mite treatment affective?" Categorical variable with the following options: "Yes"; "No" |
| **Survival_Rate_Description** | Original survey question: "What is your survival rate? Summer to Fall; Fall to unpacking / movement to yards; Two months after being taken out" Open-ended response. |
| **Summer_Climate_Zone** | Derived data field: US Climate Zone corresponding to the primary, or first listed, state location for June (Summer) 2024, extracted from "Location_June_2024." Climate Zone classifications developed by NOAA's National Centers for Environmental Information (Karl and Koss, 1984). See here: https://www.ncei.noaa.gov/access/monitoring/reference-maps/us-climate-regions |
| **survival_percent** | Derived data field: Estimated survival rate, as a percentage between 0 and 100, derived from "Survival_Rate_Description." |
| **survival_rate** | Derived data field: Estimated survival rate, as a proportion between 0 and 1, derived from "Survival_Rate_Description." |
| **loss_rate** | Derived data field: Estimated overall loss rate, as a proportion between 0 and 1, calculated as 1 - "survival_rate" |

**Supplemental Figures**

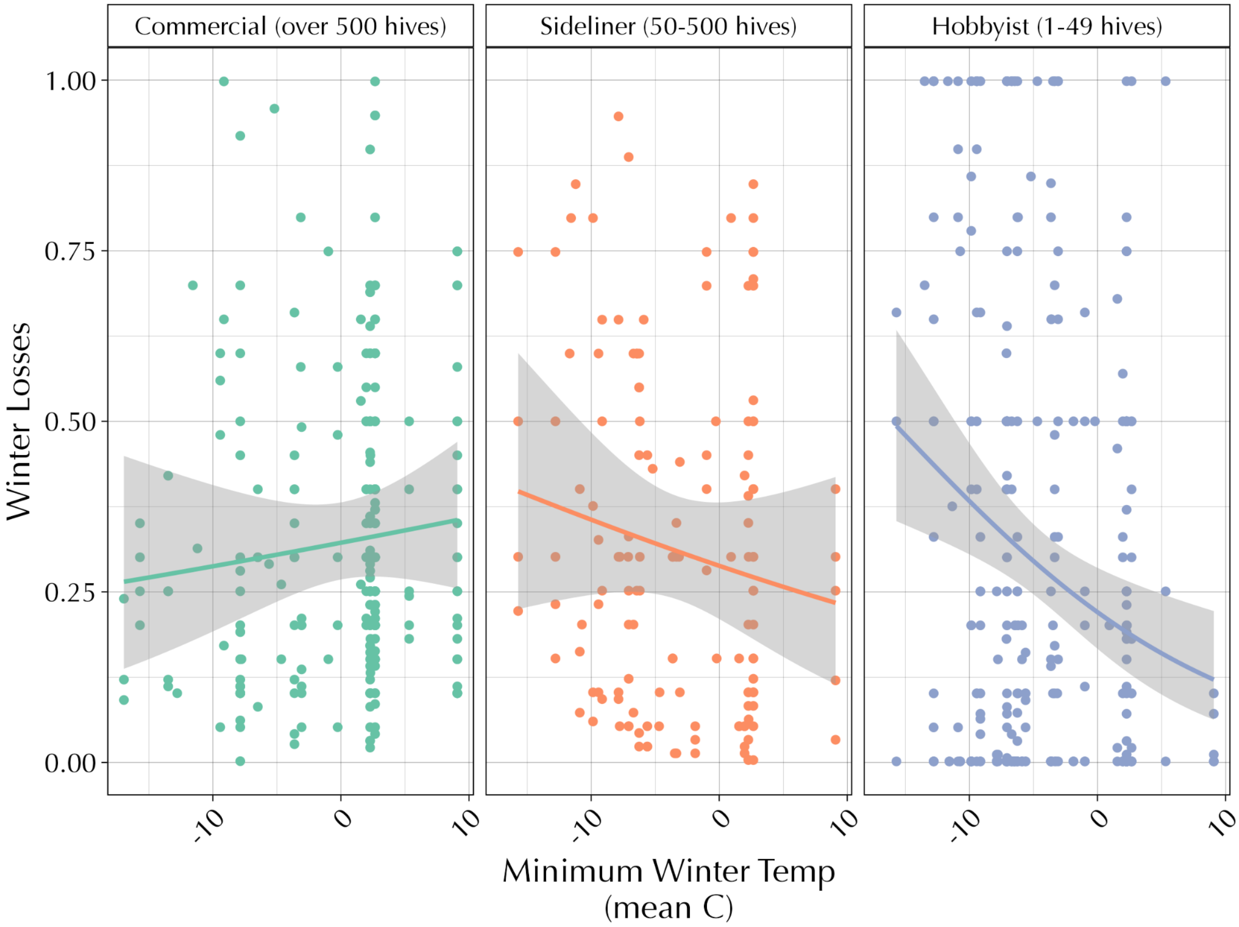


Figure S1. Winter losses as a function of mean minimum winter temperature (November – February).

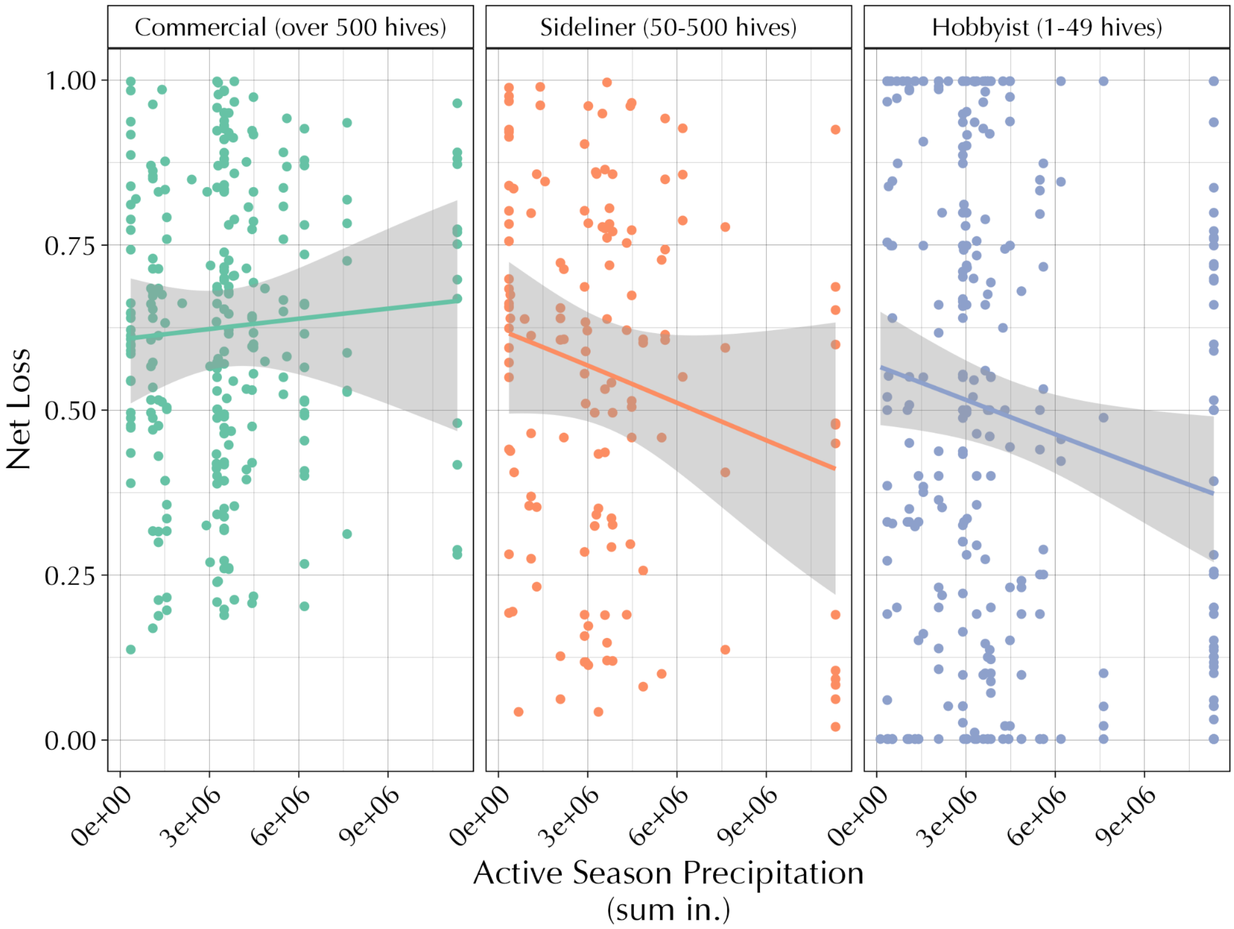


Figure S2. Net loss as a function of total (sum) active season precipitation (in., June - September).

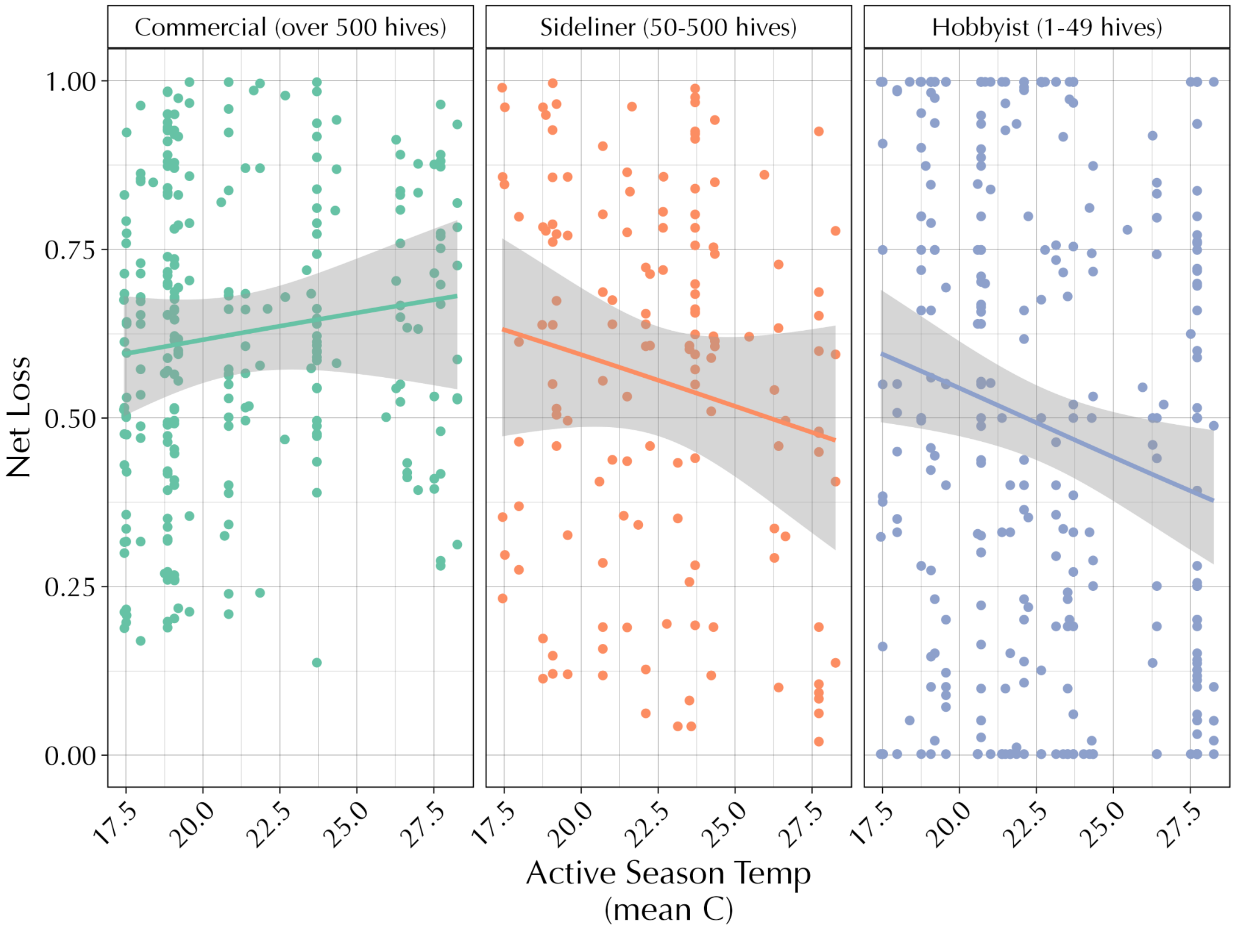


Figure S3. Net loss as a function of mean active season temperature (June - September).

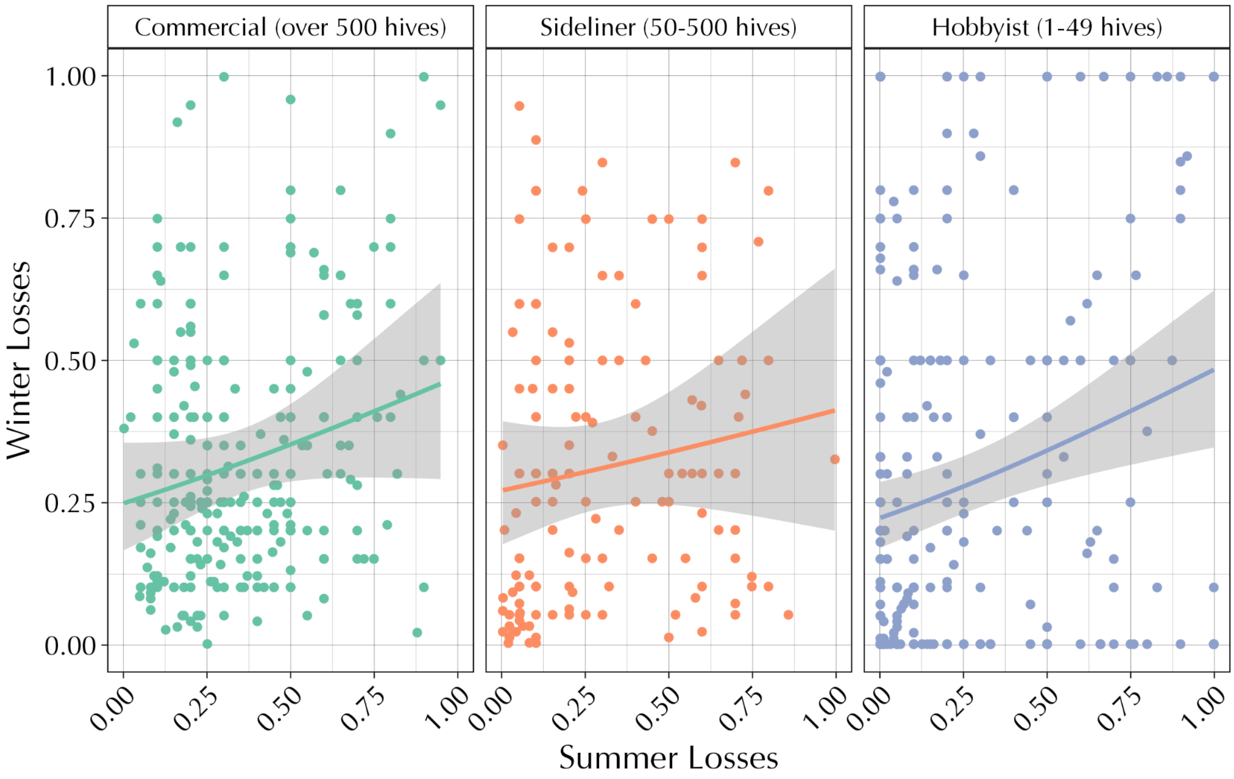


Figure S4. Winter loss as a function of summer loss.

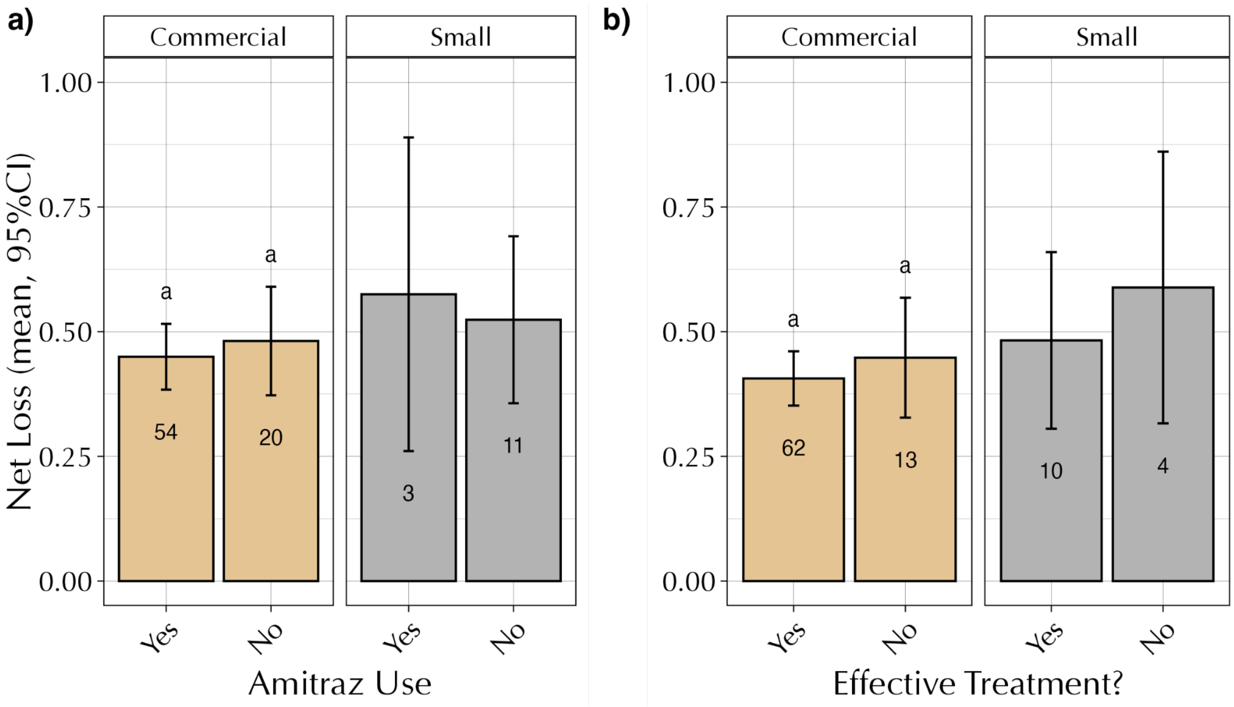

Figure S5. a) Colony loss rates by amitraz use, and b) net colony loss % for beekeepers who felt the mite treatments used were effective versus those who did not (ABF survey). Numbers under the error bars represent respondent count.

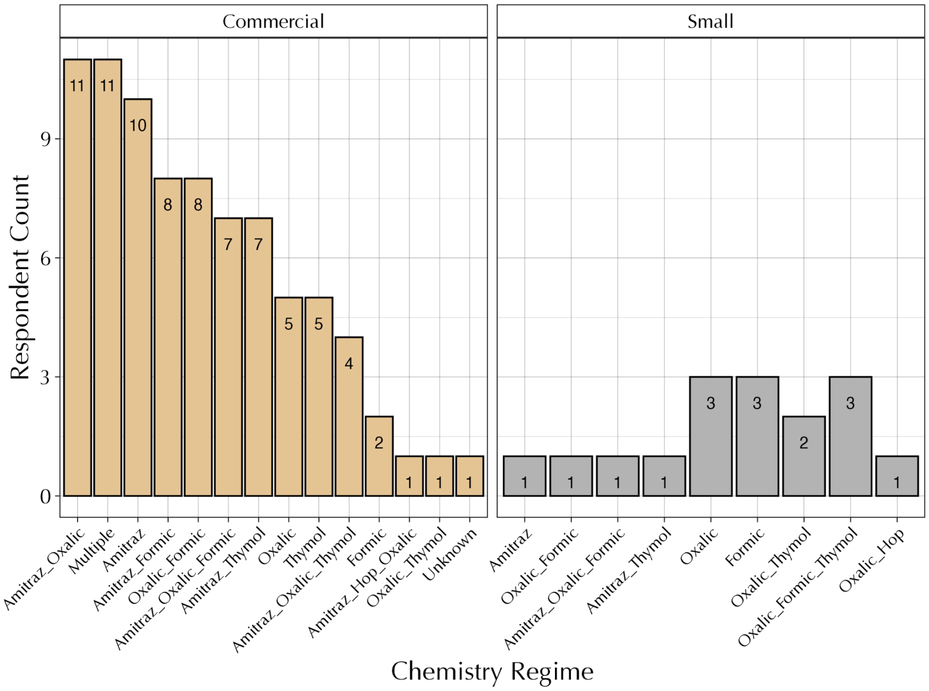


Figure S6. Described miticide use by commercial and smaller beekeepers (ABF survey).

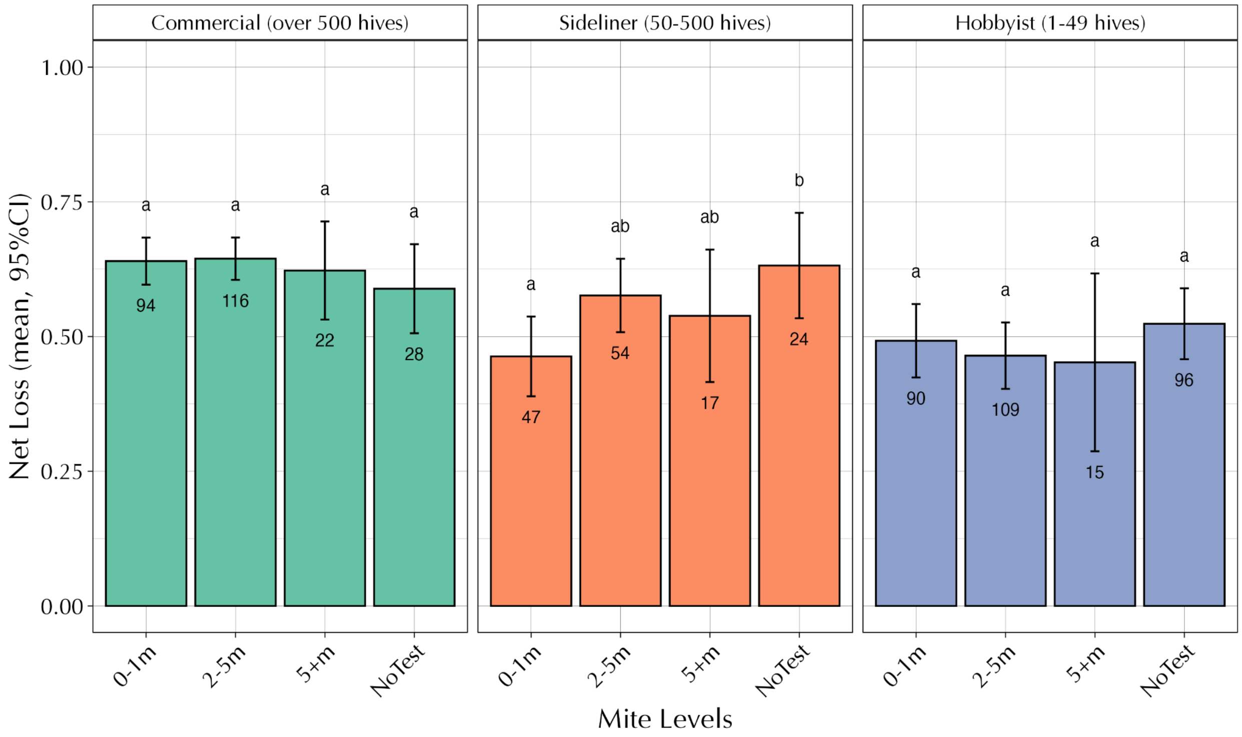


Figure S7. Colony loss by reported mite levels and beekeeper class, presented as mites/100 bees (P*Am* survey).

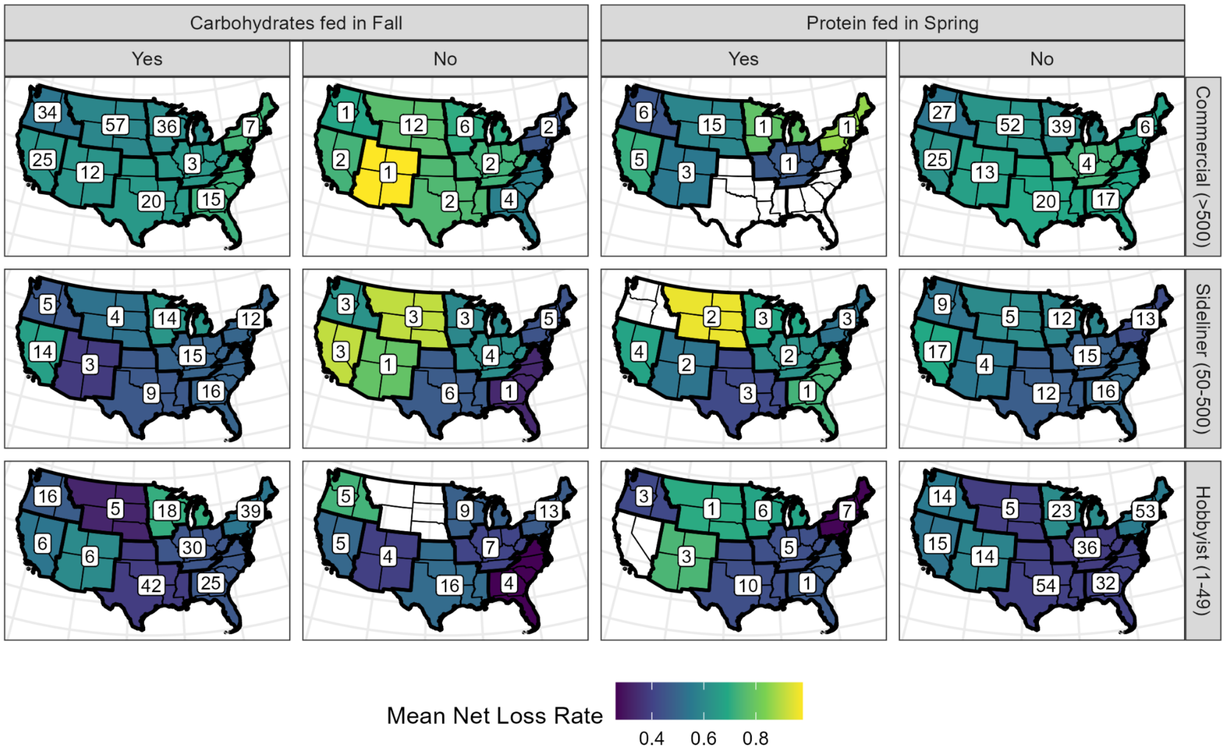


Figure S8. Mean net loss rates reported in the P*Am* beekeeper survey, with results shown for beekeepers based on whether hives were provided with supplemental carbohydrates (e.g., sugar; leftmost two columns) or supplemental protein (e.g., pollen patties; rightmost two columns). Mean net loss rates were calculated for groups of beekeepers based on the first listed summer location (binned by region according to the U.S. Climate Zone classifications delineated by NOAA; Karl and Koss 1984, link) and operation size according to the number of hives listed in the parentheses (rows). The number of beekeepers in each group are shown in white boxes superimposed on each region.

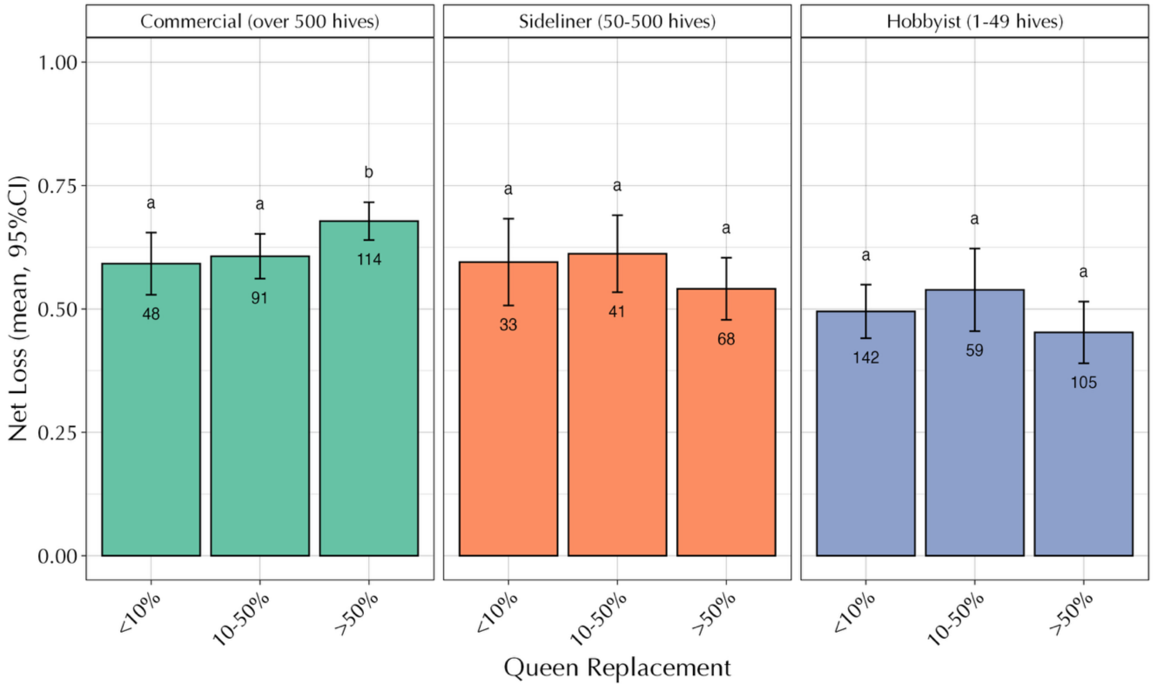


Figure S9. Colony loss as a function of queen replacement (P*Am* survey).

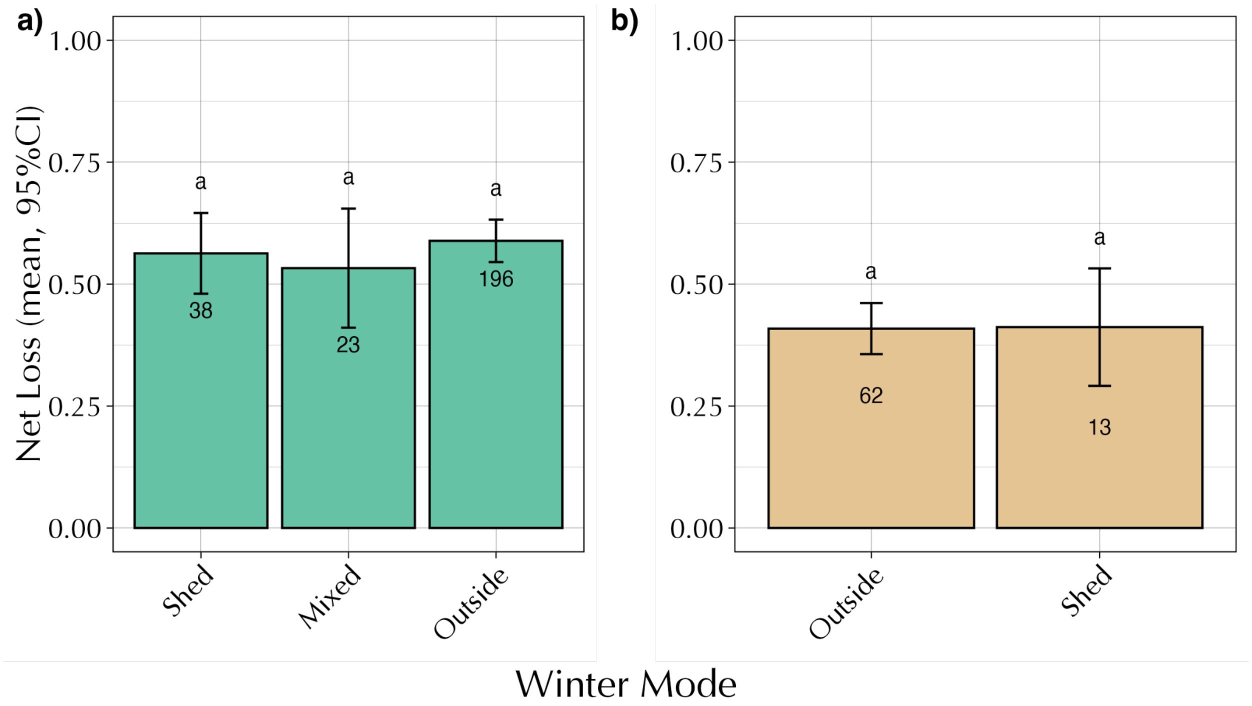


Figure S10. Colony loss as a function of winter mode (in storage sheds or outside). a) P*Am* survey of commercial beekeepers, b) ABF survey of larger (>50 colonies) beekeepers.
